# TMEM41B is an endoplasmic reticulum Ca^2+^ release channel maintaining T cell metabolic quiescence and responsiveness

**DOI:** 10.1101/2023.12.25.573330

**Authors:** Yuying Ma, Yi Wang, Xiaocui Zhao, Gang Jin, Jing Xu, Zhuoyang Li, Na Yin, Zhaobing Gao, Bingqing Xia, Min Peng

## Abstract

Naive T cells are metabolically quiescent. Here, we report an unexpected role of endoplasmic reticulum (ER) Ca^2+^ in preserving T cell metabolic quiescence. TMEM41B, an ER-resident membrane protein previously known for its crucial roles in autophagy, lipid scrabbling and viral infections, is identified as a novel type of concentration-dependent ER Ca^2+^ release channel. Ablation of TMEM41B induces ER Ca^2+^ overload, triggering the upregulation of IL-2 and IL-7 receptors in naive T cells. Consequently, this leads to increased basal signaling of the JAK-STAT, AKT-mTOR, and MAPK pathways, propelling TMEM41B-deficient naive T cells into a metabolically activated yet immunologically naive state. ER Ca^2+^ overload also downregulates CD5, a suppressor of TCR signaling, thereby reducing the activation threshold of TMEM41B-deficient T cells, resulting in attenuated tolerance and heightened T cell responses during infections. In summary, TMEM41B-mediated ER Ca^2+^ release is a pivotal determinant governing metabolic quiescence and responsiveness of naive T cells.

## INTRODUCTION

Maintaining homeostasis of naive T cells is essential for immunity and tolerance. Following maturation in the thymus, naive T cells exist in a quiescent state in the peripheral until activated by cognate antigens. The quiescence of naive T cells is not merely a passive condition but rather an actively maintained state ^1–3^. Aside from extrinsic regulations by regulatory T cells and immunosuppressive cytokines ^4,5^, the quiescence of naive T cells is intrinsically regulated at epigenetic, transcriptional, post-transcriptional and signaling transduction levels^6^. While the past decade has shed light on the critical roles of metabolic reprogramming in T cell activation, proliferation, and differentiation ^7–10^, the metabolic regulation of quiescence in resting naive T cells has been relatively understudied. Consequently, while it is established that naive T cells are metabolically quiescent^7^, the mechanisms preserving such metabolic quiescence remain poorly understood^11^.

As the central organelle for cellular metabolism, the mitochondrion has been extensively studied in T cells^12,13^, while the role of other intracellular organelles in T cell metabolism remains largely unexplored. The endoplasmic reticulum (ER) serves as the primary organelle for protein and lipid biosynthesis^14^, which also plays essential roles in Ca^2+^ signaling, particularly in lymphocytes^15^. In fact, the majority of intracellular Ca^2+^ is stored in the ER, resulting in the ER having a Ca^2+^ concentration ([Ca^2+^]_ER_) in the millimolar (mM) range, while cytosolic Ca^2+^ concentration ([Ca^2+^]_cyto_) is in the nanomolar range (nM) under steady-state^16^. This huge difference in [Ca^2+^] between ER and cytosol is established by continuous pumping of Ca^2+^ from cytosol into ER lumen against Ca^2+^ concentration gradient by the SERCA family ATPases^17^, which is balanced by ER Ca^2+^ release to cytosol. In activated T cells, T cell receptor (TCR) signaling induces rapid ER Ca^2+^ release through IP3R channels. The subsequent depletion of ER Ca^2+^ store activates the Ca^2+^ sensor STIM1/2, which, in turn, opens Ca^2+^ channels on plasma membrane, initiating Ca^2+^ influx, a phenomenon known as store-operated Ca^2+^ entry (SOCE). SOCE is crucial for T cell activation and metabolic reprogramming^18–20^. While the role of ER Ca^2+^ depletion-induced SCOE in activated T cells is well-established, whether ER Ca^2+^ plays any role in resting naive T cells was unknown. To explore how ER Ca^2+^ regulates naive T cell homeostasis, we need to understand mechanisms maintaining ER Ca^2+^ level in naive T cells. As mentioned, SERCAs continuously pump Ca^2+^ from cytosol into ER, and in activated T cells, ER Ca^2+^ can be released into cytosol via IP3R. However, the manner in which resting naive T cells release ER Ca^2+^ to the cytosol remained unclear. Over 60 years ago, passive and concentration-dependent Ca^2+^ release from the ER was described in mammalian cells^21^. Thapsigargin (TG), an inhibitor of SERCA, induces rapid ER Ca^2+^ release in virtually all metazoan cells^22,23^, indicating that ER Ca^2+^ is continuously released to cytosol under steady-state. Nevertheless, the specific Ca^2+^ channel(s) involved in this steady-state ER Ca^2+^ release in T cells remain elusive^24–26^.

Genome-wide CRISPR screens serve as powerful tools for functionally annotating uncharacterized proteins in the genome. In 2018 and 2019, two independent CRISPR screens identified TMEM41B, a poorly characterized multiple-spanning membrane protein on ER, as a critical regulator of autophagy^27,28^. Amidst the COVID-19 pandemic, while searching for viral host factors through CRISPR screening, multiple groups independently identified TMEM41B as a pan-flavivirus and pan-coronavirus (including SARS-CoV2) host factor^29–31^. Biochemically, TMEM41B exhibits phospholipid scramblases activity^32,33^, and may regulate lipid metabolism and viral infection via such activity^34–36^. Despite these insights, the biological function of TMEM41B in the ER remains enigmatic.

Here, in a genome-wide CRISPR screening aimed at identifying putative ER Ca^2+^ regulators in T cells, we unexpectedly unveil TMEM41B as a novel concentration-dependent Ca^2+^ channel to releases ER Ca^2+^, acting as a crucial mechanism to prevent ER Ca^2+^ overload. Importantly, we demonstrate a previously unappreciated but critical role of TMEM41B-mediated ER Ca^2+^ release in metabolic quiescence of naive T cells, with significant influences on T cell tolerance and responsiveness.

## RESULTS

### TMEM41B deficiency causes ER Ca^2+^ overload

To identify potential ER Ca^2+^ release channel(s) in T cells, we performed a genome-wide CRISPR screening in Jurkat T cells, using the Ca^2+^-NFAT-PD-1 axis as a readout (Figures 1A and S1A), as recently published ^37^. The rationale of this screening is that ER Ca^2+^ levels determine SOCE, which, in turn, induces PD-1 expression in T cells through a NFAT-dependent manner^19,20,38^ (Figure 1A). In this screening, PD-1 itself and known components of Ca^2+^-NFAT axis were recovered as top hits (Figure 1B). Intriguingly, two ER-resident membrane proteins, VMP1 and TMEM41B, were also identified as top hits (Figure 1B). We have shown that VMP1 regulates ER Ca^2+^ and naive T cell survival^37^. In this study, we focus on the role of TMEM41B in ER Ca^2+^ regulation and T cell homeostasis.

**Figure 1.**
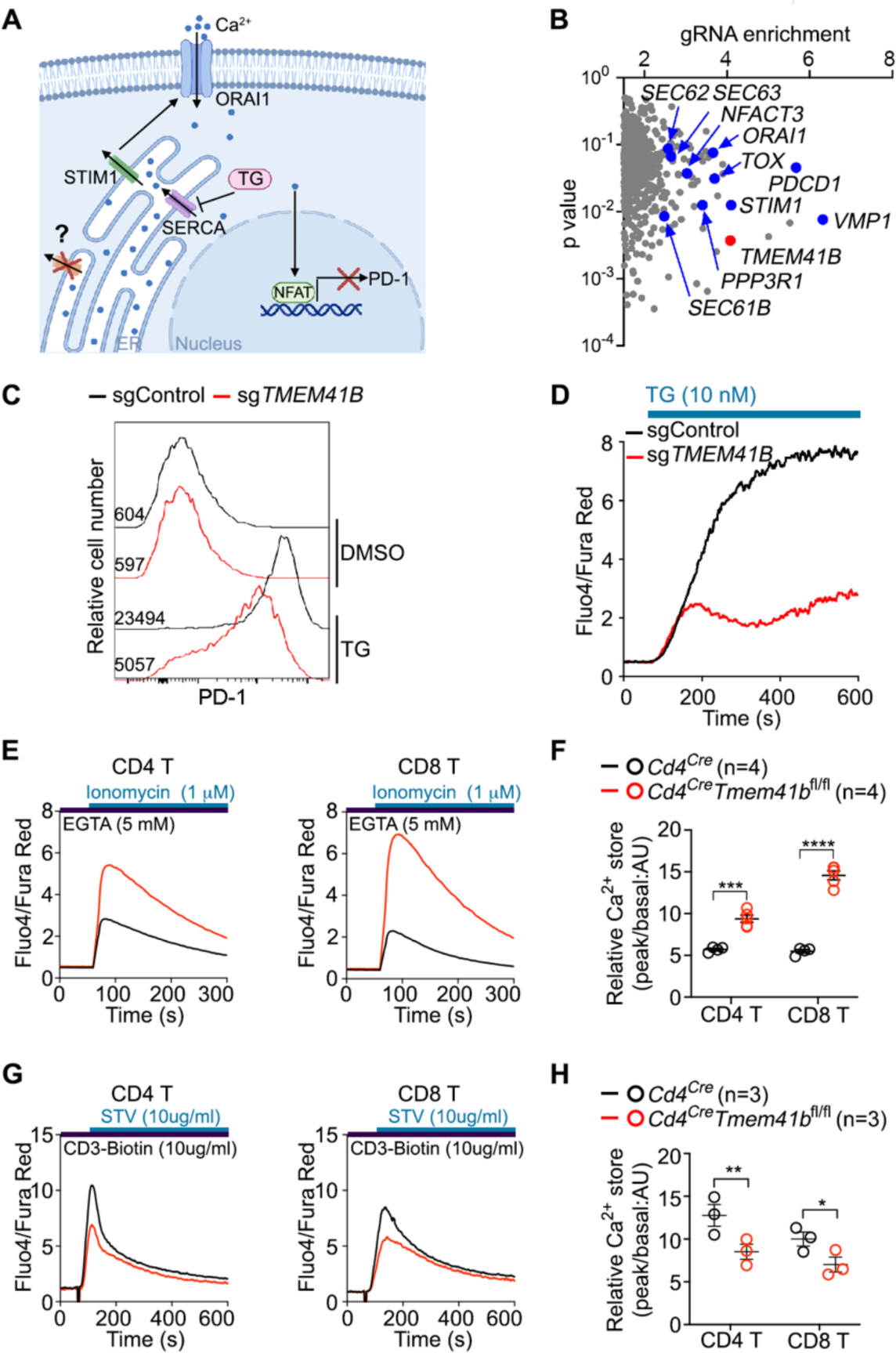
TMEM41B deficiency causes ER Ca^2+^ overload. (A) A cartoon illustrating store-operated Ca^2+^ entry (SOCE) and thapsigargin (TG)-Ca^2+^-NFAT-PD-1 signaling axis in T cells. (B) Screening result from Jurkat cells. Selected hits are labelled. (C) Flow cytometry analysis of PD-1 expression by Jurkat cells (Cas9^+^) expressing indicated sgRNAs upon treatment with DMSO or TG (10 nM) for 48 h. Representative plots from 3 independent experiments are shown. (D) Flow cytometry analysis of TG-induced Ca^2+^ influx in Jurkat cells (Cas9^+^) expressing indicated sgRNAs. Representative plots from 3 independent experiments are shown. (E and F) Flow cytometry analysis of ER Ca^2+^ store in T cells with indicated genotypes. Representative plots (E) and statistics (F) are shown. (n = 4 mice) (G and H) Flow cytometry analysis of αCD3-induced Ca^2+^ influx in T cells with indicated genotypes. Representative plots (G) and statistics (H) are shown. (n = 3 mice) * p < 0.05, ** p < 0.01, *** p < 0.001, **** p< 0.0001 by two-tailed unpaired t-test (F and H).

We confirmed that thapsigargin-induced PD-1 upregulation was diminished in TMEM41B-deficient Jurkat T cells (Figure 1C). Additionally, thapsigargin-induced Ca^2+^ influx was attenuated in TMEM41B-deficient Jurkat T cells (Figure 1D). This reduction in PD-1 induction and Ca^2+^ influx upon thapsigargin treatment was further validated in TMEM41B-deficient primary mouse T cells (Figures S1B - S1D). In a standard SOCE assay, we observed blunted SOCE in TMEM41B-deficient HEK293T cells, along with increased ER Ca^2+^ levels in these cells (Figures S1E - S1G), a phenomenon further valiadted in two monoclonal TMEM41B knockout HEK293T clones (Figures S1H - S1J). To directly monitor ER Ca^2+^ level, we utilized a HEK293T cell line that stably expressed the ER Ca^2+^ sensor G-CEPIA1er^39^. In HEK293T-G-CEPIA1er cells, TMEM41B deficiency significantly increased ER Ca^2+^ level, both under steady state and after thapsigargin treatment (Figures S1K - S1M). These findings collectively demonstrate that TMEM41B promotes ER Ca^2+^ release.

ER Ca^2+^ level was significantly increased in TMEM41B-deficient naive T cells compared to control naive T cells (Figures 1E and 1F). Ca^2+^ influx induced by CD3 cross-linking was partially reduced in TMEM41B-deficent T cells (Figures 1G and 1H), consistent with increased ER Ca^2+^ levels in these cells (Figures 1E and 1F), as SOCE is determined by ER Ca^2+^ levels.

Together, these data establish that TMEM41B deficiency leads to ER Ca^2+^ overload in T cells, partially impairing SOCE.

### TMEM41B mediates Ca^2+^ transport across membranes

To assess whether TMEM41B is sufficient for ER Ca^2+^ release, we overexpressed TMEM41B in HEK293T cells (Figure 2A). The overexpression of TMEM41B resulted in near-complete depletion of ER Ca^2+^ in HEK293T cells (Figures 2B and 2C). In HEK293T-G-CEPIA1er cells, the steady-state ER Ca^2+^ levels were almost completely depleted upon TMEM41B overexpression, and minimal ER Ca^2+^ release was observed after thapsigargin treatment (Figures 2D and 2E). These data demonstrate that Ca^2+^ in the ER is almost entirely released upon TMEM41B overexpression.

**Figure 2.**
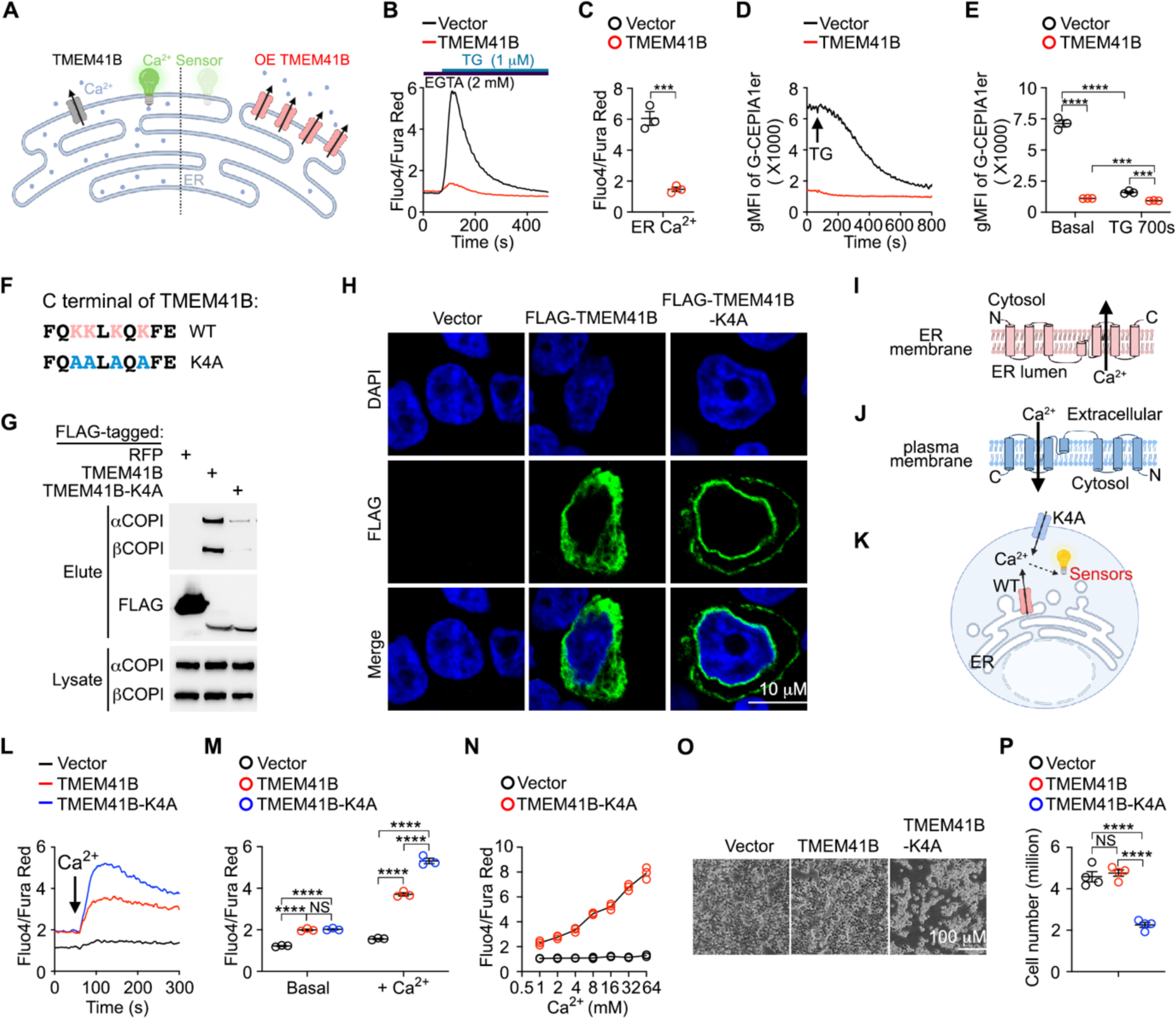
TMEM41B mediates Ca^2+^ transport across membranes. (A) A cartoon illustrating TMEM41B overexpression and its potential influence on ER Ca^2+^. (B and C) Flow cytometry analysis of ER Ca^2+^ in HEK293T cells transfected with indicated plasmids. Representative plots (B) and statistics (C) are shown. (n = 3 independent experiments). (D and E) Flow cytometry analysis of ER Ca^2+^ in HEK293T-G-CEPIA1er cells transfected with indicated plasmids before and after TG (1 μM) treatment. Representative plots (D) and statistics (E) are shown. (n = 3 independent experiments). (F) C-terminal amino acid sequence of TMEM41B. Lysines in Golgi-to-ER retrieve motif are labeled in red and mutated alanines are labeled in blue. (G) The interaction between RFP, TMEM41B or TMEM41B-K4A with COPI was examined by coimmunoprecipitation. The first lane was used in our previous study^37^. (H) Immunofluorescence examination of subcellular localization of FLAG-tagged (N-terminal) TMEM41B and TMEM41B-K4A. (I and J) Putative topology of TMEM41B on ER (I) and TMEM41B-K4A on plasma membrane (J). (K) Putative Ca^2+^ transport by TMEM41B on ER and TMEM41B-K4A on plasma membrane. (L and M) Flow cytometry analysis of Ca^2+^ influx in HEK293T cells transfected with indicated plasmids. Representative plots (L) and statistics (M) are shown. (n = 3 independent experiments). (N) Flow cytometry analysis of Ca^2+^ influx in HEK293T cells transfected with indicated plasmids. (n = 3 independent experiments). (O and P) Survival of HEK293T cells transfected with indicated plasmids (48 h post-transfection). Representative images (O) and statistics (P) are shown. (n = 4 independent experiments). *** p < 0.001, **** p< 0.0001, NS, not significant by two-tailed unpaired t-test (C), two-way ANOVA (E, M), one-way ANOVA (P).

Given that TMEM41B is an ER-resident membrane protein^27,28^, it is technically challenging to measure Ca^2+^ transport across the ER membrane. We thus redirected TMEM41B to plasma membrane using a strategy that we put VMP1 on plasma membrane in our recent study^37^. TMEM41B possesses a lysine-rich Golgi-to-ER retrieve motif at its C-terminus (Figure 2F), which interacts with COPI complex facilitating the retro-transport of membrane proteins from the trans-Golgi back to ER^40^. When lysine residues within this Golgi-to-ER retrieval motif were mutated to alanine (TMEM41B-K4A), the resulting mutant exhibited reduced COPI binding compared to its wild-type counterpart (Figure 2G). Unlike wild-type TMEM41B exhibiting typical ER localization, TMEM41B-K4A showed plasma membrane localization in addition to the ER (Figure 2H).

According to topology of membrane proteins, lumen side of TMEM41B should face extracellularly when expressed on plasma membrane (Figures 2I and 2J). Since Ca^2+^ concentrations are high in extracellular space (∼ mM range) and low in cytosol (∼ nM range), plasma membrane-targeted TMEM41B-K4A should induce Ca^2+^ influx if it is capable of channeling Ca^2+^ in the presence of electrochemical gradients of Ca^2+^ (Figure 2K). In empty vector-transfected HEK293T cells, Ca^2+^ supplementation induced neglectable Ca^2+^ influx (Figures 2L and 2M), as expected since mammalian cells do not express concentration-dependent Ca^2+^ channels on the cell surface^41^. In cells overexpressing wild-type TMEM41B, Ca^2+^ supplementation induced a small Ca^2+^ influx, while robust Ca^2+^ influx was observed in cells overexpressing the plasma membrane-targeted TMEM41B-K4A (Figures 2L and 2M). The Ca^2+^ influx in cells overexpressing TMEM41B-K4A was concentration-dependent (Figure 2N). Prolonged Ca^2+^ influx is known to trigger cell death^22,23^. Indeed, HEK293T cells overexpressing plasma membrane-targeted TMEM41B-K4A exhibited increased cell death compared with control cells (Figures 2O and 2P).

To investigate whether the Ca^2+^ influx observed in HEK293T cells overexpressing TMEM41B or TMEM41B-K4A was SOCE, we repeated these experiments in STIM1- or ORAI1-deficient HEK293T cells incapable of inducing SCOE^37^. Ca^2+^ influx induced by TMEM41B overexpression was abolished in STIM1- or ORAI1-deficient HEK293T cells, while TMEM41B-K4A-induced Ca^2+^ influx remained unaffected (Figures S2A - S2C), demonstrating that Ca^2+^ influx induced by TMEM41B overexpression is SOCE (due to its depletion of ER Ca^2+^), while TMEM41B-K4A induced Ca^2+^ influx is not SOCE.

Together, these data demonstrate that TMEM41B is able to meidate Ca^2+^ transport across membranes, either by acting as a Ca^2+^ channel itself or by facilitating the opening Ca^2+^ channel(s) other than ORAI1.

### TMEM41B forms a Ca^2+^ channel

To investigate whether TMEM41B functions as a Ca^2+^ channel, we conducted single-channel electrophysiology using purified recombinant TMEM41B (Figures S2D – S2F). Although the molecular weight of TMEM41B was 25 kDa on denatured gels (Figures S2E and S2F), TMEM41B appeared significantly larger than the monomeric form on native gels (Figures S2G and S2H). These data suggest that purified TMEM41B exists as oligomers, a typical feature of ion channels.

Single-channel currents of TMEM41B were recorded using planar lipid bilayer experiments (Figure 3A). We successfully recorded single-channel currents of TMEM41B with a conductance of 37.18 ± 7.33 pS (mean ± SEM) (Figures 3B - 3D). As a control, the elution buffer (containing FLAG peptide) did not exhibit detectable currents (Figure 3B). In an asymmetric 50: 500 mM KCl solution, TMEM41B displayed a reverse potential of 52.77 mV, resulting in an estimated P_K+_/P_Cl_- of 31.77, indicating that TMEM41B is a cation-selective channel (Figures 3C and 3D).

**Figure 3.**
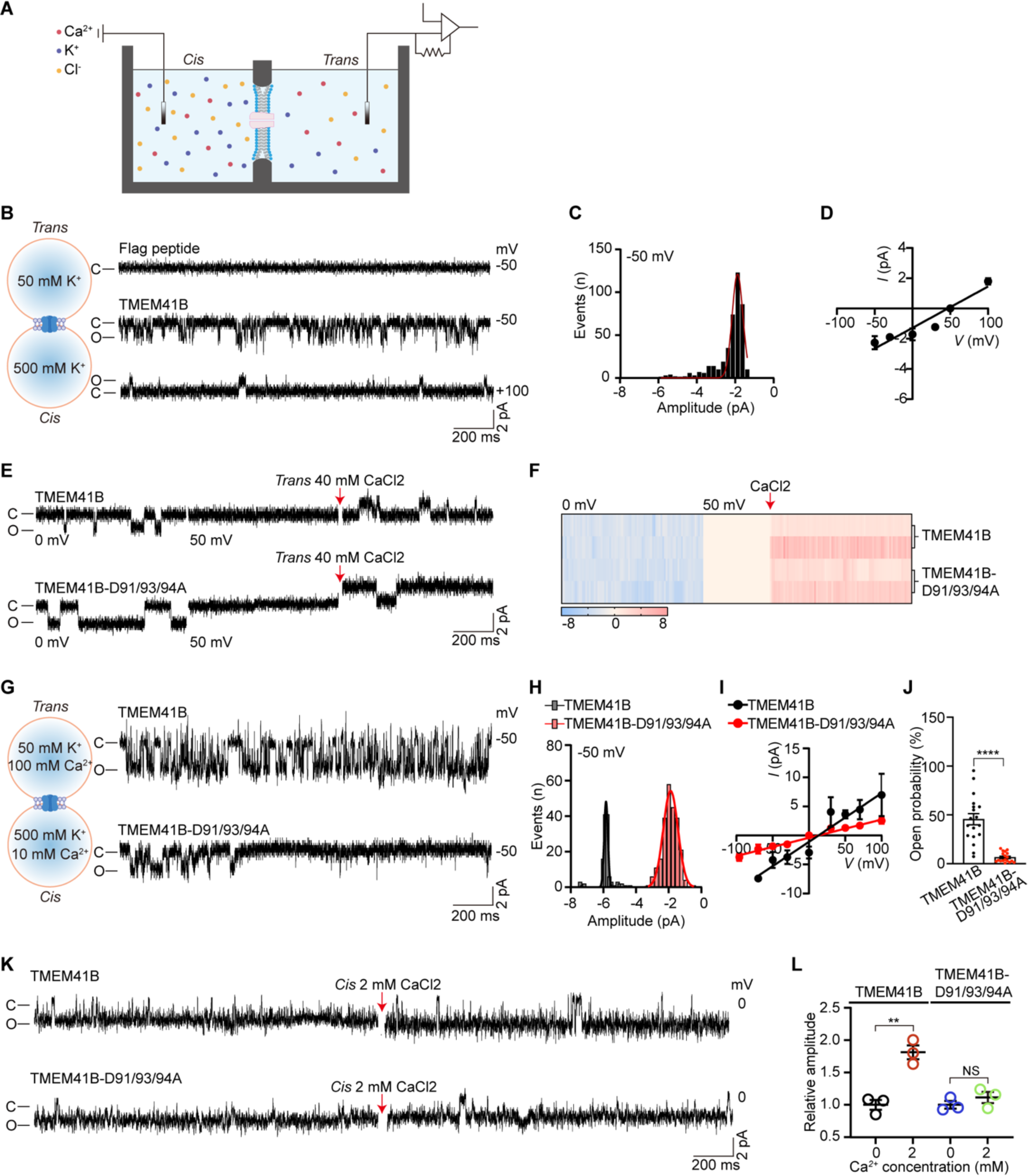
TMEM41B forms a Ca^2+^ channel. (A) A carton of planar lipid bilayer work station (BLM Workstation). (B) Left, solution conditions in *Cis* and *Trans* side. Right, representative single channel currents of TMEM41B or Flag peptide at indicated voltage. “C” means closed; “O” means open. (C) All-point current histograms for the trace in (B). (D) I-V curve of TMEM41B in solution of (B). (E) Representative single channel currents of TMEM41B and TMEM41B-D91/93/94A at indicated voltage. Red arrow indicates that the CaCl_2_ has been added. (F) Heat map of amplitude in (E). (G) Left, solution conditions in *Cis* and *Trans* side. Right, representative single channel currents of TMEM41B and TMEM41B-D91/93/94A at -50 mV. (H) All-point current histograms for the trace in (G). (I) I-V curve of TMEM41B and TMEM41B-D91/93/94A in solution of (G). (J) Open probability of TMEM41B and TMEM41B-D91/93/94A in solution of (G). (n = 18 independent experiments) (K) Representative single channel currents of TMEM41B and TMEM41B-D91/93/94A at indicated voltage. Red arrow indicates that the CaCl_2_ has been added. (L) Relative amplitude of TMEM41B and TMEM41B-D91/93/94A in (K). (n = 3 independent experiments) ** p < 0.01, **** p < 0.0001, NS, not significant by two-tailed unpaired t-test (J and L).

To explore the permeability of TMEM41B to Ca^2+^, Ca^2+^ currents were analyzed. The membrane potential was adjusted to the K^+^ equilibrium potential (50 mV) to eliminate K^+^ currents, after which Ca^2+^ was added for recording. Inward step-like signals were observed when 40 mM (final concentration) Ca^2+^ was added to the *Trans* side (Figures 3E and 3F). Additionally, addition of asymmetric Ca^2+^ (100: 10 mM) into the bath solution caused the reversal potential of TMEM41B to shift from 52.77 mV to 13.04 mV (Figures 3G - 3I). This substantial shift of reversal potential suggests that TMEM41B is selective for Ca^2+^. Importantly, the conductance of TMEM41B increased to 83.7±10.85 pS after adding CaCl_2_ in the bath solution, indicating that TMEM41B is a Ca^2+^-regulated channel (Figure 3I).

To identify key residues of TMEM41B invovled in Ca^2+^ channelling, we individually mutated all negatively charged amino acids (aspartic acid and glutamic acid) in TMEM41B facing the ER lumen side. In a rescue system, where TMEM41B and mutants were re-expressed in TMEM41B-deficient primary T cells to reverse ER Ca^2+^ overload caused by TMEM41B deficiency, we found that none of these single amino acid mutants exhibited loss of function in our reconstitution assay (data not shown). However, when aspartate 91/93/94 (D91/93/94) were mutated to alanine (D91/93/94A) simultaneously, a partial loss of function in TMEM41B-mediated ER Ca^2+^ release was observed (Figures S2I – S2K).

Subsequently, the D91/93/94A mutant was purified and recorded accordingly. Electrophysiological results revealed that the D91/93/94A mutant could form a cation channel similar to wild-type TMEM41B (Erev = 46.96 mV; P_K+_/P_Cl-_ = 15.38) (Figures S2L – S2N). While the D91/93/94A mutant remained permeable to Ca^2+^ (Erev = 12.81 mV; P_K+_/P_Ca2+_ = 1.73), its conductance reduced from 83.70 ± 10.85 pS to 31.44 ± 2.23 in the K^+^/Ca^2+^ mixture solution (Figures 3G - 3I). Moreover, the open probability of the D91/93/94A mutant significantly decreased compared with wild-type TMEM41B in the mixture solution (Figure 3J). Thus, D91/93/94 are critical for Ca^2+^ permeability of the TMEM41B channel.

Finally, the influence of Ca^2+^ on TMEM41B channel was evaluated using titration strategy on the same channel. After channels were incorporated into membranes, Ca^2+^ (2 mM CaCl_2_) was titrated into the *Cis* side. As expected, the addition of Ca^2+^ significantly increased the amplitude of wild-type channels, while Ca^2+^ failed to increase the channel activity of the D91/93/94A mutant (Figures 3K and 3L). In contrast, the open probability (P_open_) of wild-type and D91/93/94A mutant of TMEM41B was comparable, and also not affected by the addition of Ca^2+^ (Figure S3O). These results support the notion that D91/93/94 are key residues determining the Ca^2+^ preference and Ca^2+^ permeability of the TMEM41B channel.

With our cell based assay demonstrating that TMEM41B induces Ca^2+^ influx in a concentration-dependent manner (Figure 2N), these data collectively demonstrate that TMEM41B forms a Ca^2+^ channel, and the activity of this channel depends on Ca^2+^ concentration.

### TMEM41B maintains metabolic quiescence of naive T cells

Having established TMEM41B as an ER Ca^2+^ release channel, we proceeded to investigate its role in T cells in vivo. We generated T cell-specific *Tmem41b* knockout mice (*Cd4^Cre^Tmem41b*^fl/fl^) through the crossing of *Tmem41b* floxed mice with *Cd4^Cre^* transgenic mice (Figure S3A). *Tmem41b* knockout was confirmed through PCR analysis (Figures S3B and S3C). Deletion of *Tmem41b* with *Cd4^Cre^* did not affect T cell development in thymus (Figures S3D – S3F). While there was a slight reduction in mature CD8 T cells in the secondary lymphoid organs of *Cd4^Cre^Tmem41b*^fl/fl^ mice (Figure S3G), apoptosis of TMEM41B*-*deficient CD8 T cells was unchanged (Figures S3H and S3I). The observed decrease in CD8 T cell number is likely attributed to heightened ER stress in these cells (Figures S3J and S3K). However, unlike VMP1*-*deficient T cells, TMEM41B*-* deficient T cells did not exhibit mitochondrial Ca^2+^ overload (Figures S3L and S3M), which caused massive apoptosis of VMP1*-*deficient T cells^37^. Thus, TMEM41B plays a distinct role in T cells compared to VMP1.

Flow cytometry analysis revealed that TMEM41B-deficient T cells and wild-type T cells were indistinguishable in terms of their T cell activation state (Figures 4A, 4B, S4A and S4B). However, TMEM41B-deficient naive T cells exhibited larger sizes compared to control naive T cells (Figures 4C and 4D), suggesting an increase in anabolic activities.

**Figure 4.**
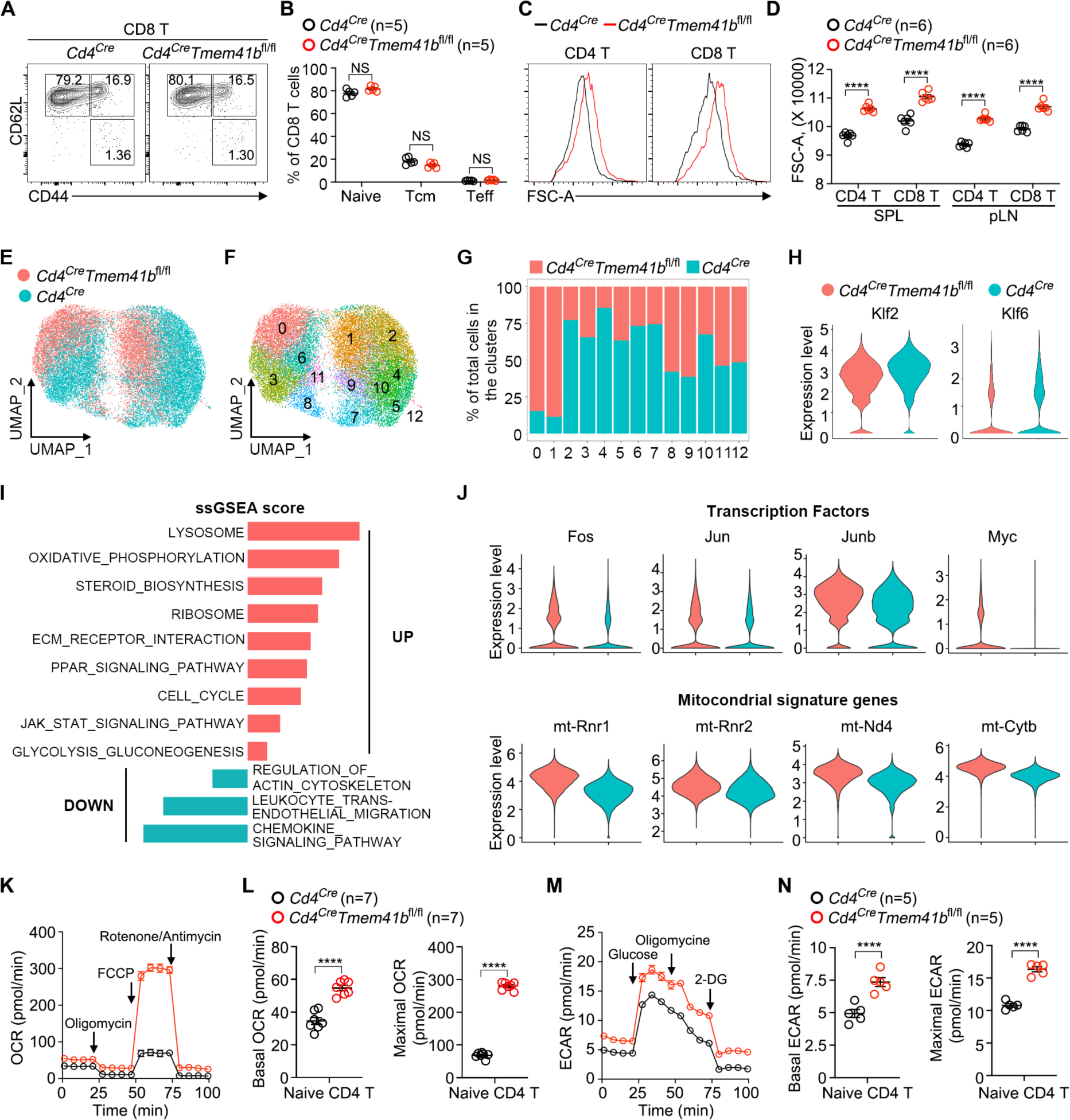
TMEM41B maintains metabolic quiescence of naive T cells. (A and B) Flow cytometry analysis of activation status (CD44 vs CD62L) of CD8 T cells from peripheral lymph nodes (pLN) of control (*Cd4*^Cre^) and TMEM41B-deficient (*Cd4*^Cre^*Tmem41b*^fl/fl^) mice (6 ∼ 8-week-old). Representative plots (A) and statistics (B) are shown. (n = 5 mice). (C and D) Flow cytometry analysis of the size of control and TMEM41B-deficient T cells from spleen and pLN. Representative plots (C) and statistics (D) are shown. (n = 6 mice). (E) A merged uniform manifold approximation and projection (UMAP) plot of pooled naive T cells from control and TMEM41B-deficient mice. (F) A UMAP plot of naive T cells from control and TMEM41B-deficient mice colored according to cluster classification. (G) The percentages of control and TMEM41B-deficeint naive T cells in each cluster. (H) Villon plots of gene expression in control and TMEM41B-deficeint naive T cells. (I) Gene-set enrichment analysis of differentially expressed genes by ssGSEA method. (J) Villon plots of gene expression in control and TMEM41B-deficeint naive T cells. (K and L) Oxygen consumption rate (OCR) of control and TMEM41B-deficeint naive CD4 T cells were measured with a Mito stress test kit. Plots (K) and calculated basal and maximal OCR (L) are shown. (n = 7 independent experiments). (M and N) Extracellular acidification rate (ECAR) of control and TMEM41B-deficeint naive CD4 T cells were measured with a Glycolysis stress test kit. Plots (M) and calculated basal and maximal ECAR (N) are shown. (n = 5 independent experiments) **** p< 0.0001, NS, not significant by two-tailed unpaired t-test (B, D, L and N).

To elucidate the underlying mechanisms, we conducted single-cell RNA sequencing (scRNA-seq) of naive T cells (CD62L^+^CD44^-^CD25^-^) isolated from control and TMEM41B-deficient mice. In concordance with the flow cytometry analysis (Figures 4A, 4B, S4A and S4B), both wild-type and TMEM41B-deficient naive T cells did not express *Cd44*, but highly expressed *Sell* (CD62L) (Figure S4C). Strikingly, the UMAP analysis revealed largely non-overlapping clusters for control and TMEM41B-deficient naive T cells (Figures 4E, S4D and S4E), indicating the existence of distinct cellular states. While TMEM41B-deficient naive T cells predominantly occupied clusters 0 and 1 (C0 and C1), control naive T cells were notably excluded from these two clusters (Figures 4F, 4G, S4D and S4E). Despite being phenotypically naive (CD62L^+^CD44^-^) (Figures 4A, 4B, S4A - S4C), TMEM41B-deficient naive T cells displayed downregulation of transcription factors associated with T cell quiescence, including *Klf2* and *Klf6* ^42^ (Figure 4H). Importantly, pathways associated with mitochondrial metabolism, glycolysis, ribosome biogenesis, PPAR and JAK-STAT signaling were enriched in TMEM41B-deficient naive T cells (Figures 4I, S4F - S4H). Specifically, genes conventionally associated with T cell activation such as *Fos*, *Jun*, *Junb* and *Myc*, along with mitochondrial genes *mtRnr1*, *mtRnr2*, *mt-Nd4*, and *mt-Cytb*, were upregulated in TMEM41B-deficient naive T cells (Figure 4J). These transcriptional changes suggest that, although TMEM41B-deficient naive T cells are not activated immunologically, they are in a metabolically active state. Indeed, metabolic assays revealed a significant increase in both oxygen consumption rate (OCR) and extracellular acidification rate (EACR) in TMEM41B-deficient naive T cells (Figures 4K - 4N), underscoring their heightened metabolic activity in a non-activated state. Consistent with increased OCR, TMEM41B-deficient T cells displayed increased mitochondrial mass and mitochondrial membrane potential (Figures S4I - S4M), as well as increased reactive oxygen species (ROS) (Figures S4N and S4O).

Collectively, these findings demonstrate that TMEM41B deficiency propels naive T cells into a metabolically activated yet immunologically naive state, a phenomenon not reported previously (See **DUSCUSSION**).

### TMEM41B represses IL-2 and IL-7 signaling in naive T cells via Ca^2+^ channel activity

To unravel the mechanisms underlying the metabolic activation, but not immunological activation, of TMEM41B-deficient naive T cells, we screened receptors involved in signaling and metabolism, including TCR, CD3, CD28, ICOS, IL-2Rα (CD25), IL-2Rβ (CD122), IL-2Rγ (CD132) and IL-7R (CD127). Consistent with the immunologically naive state of TMEM41B-deficient T cells (Figures 4A, 4B, S4A - S4C), the expression of receptors for T cell activation such as TCR, CD3, CD28 and ICOS were largely comparable between wild-type and TMEM41B-deficient naive T cells (Figures 5A and S5A). In contrast, receptors associated with promoting metabolism but incapable of inducing T cell activation independently, comprising all components of the IL-2 receptor (CD25, CD122, and CD132) and IL-7R, were consistently elevated in TMEM41B-deficient naive T cells compared to control naive T cells (Figures 5A and S5A). Subsequent examination of downstream signaling events from IL-2 and IL-7 receptors, including JAK-STAT, AKT-mTOR, and MAPK pathways, revealed increased phosphorylation of AKT, p70S6K, S6, STAT5, and ERK in TMEM41B-deficient naive T cells relative to control naive T cells (Figure 5B and S5B), indicating enhanced signaling from IL-2 and/or IL-7 receptors in TMEM41B-deficient naive T cells in the absence of T cell activation.

**Figure 5.**
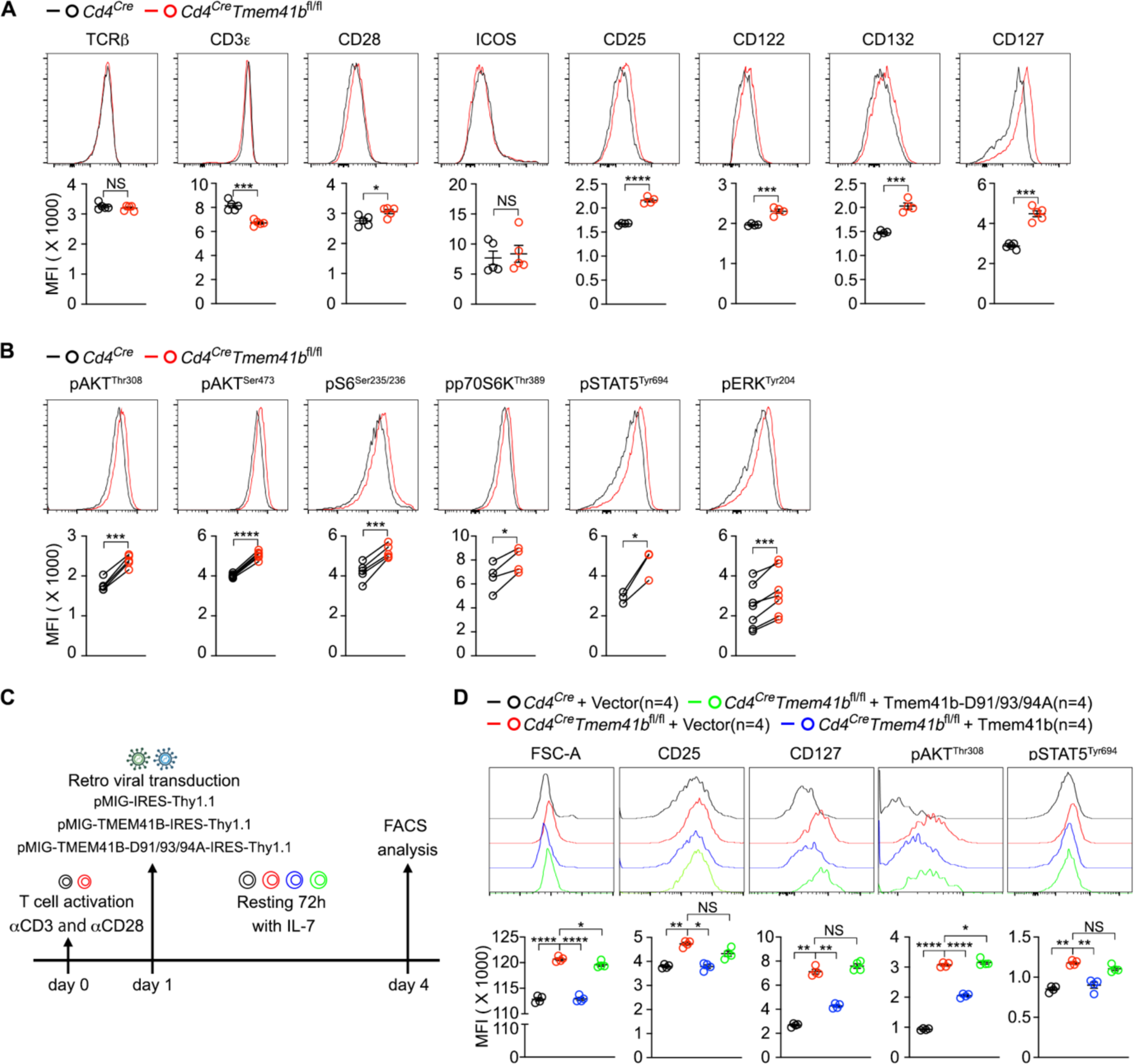
TMEM41B represses basal IL-2 and IL-7 receptor signaling in naive T cells via Ca^2+^ channel activity. (A) Flow cytometry analysis of the expression of indicated proteins on naive CD4 T cells from peripheral lymph nodes (pLN) of control (*Cd4*^Cre^) and TMEM41B-deficient (*Cd4*^Cre^*Tmem41b*^fl/fl^) mice. Representative plots and statistics are shown. (n = 4 or 5 mice). (B) Flow cytometry analysis of the phosphorylation of indicated proteins in naive CD4 T cells from pLN of control and TMEM41B-deficient mice. Representative plots and statistis are shown. (n = 3 ∼ 8 mice). (C) Experimental design for the rescue of phenotypes in TMEM41B-deficient T cells by wild-type and D91/93/94A mutant of TMEM41B. (D) Activated CD4 T cells were transduced with indicated retroviral constructs. Cell size, CD25/CD127 expression and phosphorylation of AKT/STAT5 were measured by flow cytometry 72 hours post-transduction. Representative plots and statistics are shown. (n = 4 independent experiments). * p< 0.05, ** p< 0.01, *** p< 0.001, **** p< 0.0001, NS, not significant by two-tailed unpaired t-test (A), two-tailed paired t-test (B) or one-way ANOVA (C).

To elucidate the relationship between ER Ca^2+^ overload and increased IL-2/IL-7 receptor signaling in TMEM41B-deficient T cells, we performed a rescue experiment using wild-type and D91/93/94A mutant TMEM41B, which exhibited reduced Ca^2+^ channel activity (Figure 3). In these experiments, control and TMEM41B-deficient T cells were first activated for retroviral transduction of TMEM41B or its D91/93/94A mutant, followed by a 3-day rest with IL-7 treatment to allow cells to enter a relatively quiescent state (Figure 5C). Under this condition, the upregulation of CD25 and CD127, increased AKT and STAT5 signaling, as well as enlarged cell size of TMEM41B-deficient T cells, were all reversed by the wild-type but not D91/93/94A mutant of TMEM41B (Figures 5D and S5C), indicating that all these changes are associated with ER Ca^2+^ dysregulation.

Cell size is largely determined by protein content, and the mTORC1 pathway plays critical roles in the biosynthesis and metabolism of T cells^43^. As mTORC1 is active in TMEM41B-deficient T cells (Figures 5C and S5B), we investigated whether inhibition of mTORC1 could rescue metabolic phenotypes of these cells. We crossed *Cd4^Cre^Tmem41b*^fl/fl^ mice with *Rptor* flox mice to deplete RAPTOR (Figure S6A), the defining component of the mTORC1^44^. RAPTOR deficiency largely reversed the enlarged size of TMEM41B-deficient T cells (Figures S6B and S6C). Increased OCR and ECAR in TMEM41B-deficient T cells were also partially mitigated by RAPTOR deficiency (Figures S6D - S6G). These findings demonstrate that heightened mTORC1 signaling contributes to the metabolic activation of TMEM41B-deficient T cells. However, the incomplete rescue by RAPTOR deficiency suggests that other pathways, such as STAT5 or ERK, might also contribute to the metabolic activation of TMEM41B-deficient T cells.

Collectively, these data demonstrate that ER Ca^2+^ overload in TMEM41B-deficient T cells, through yet unidentified mechanism(s), upregulates IL-2/IL-7 receptors, which leads to constitutive JAK-STAT, AKT-mTOR and MAPK signaling, ultimately resulting in the metabolic activation of TMEM41B-deficient naive T cells.

### TMEM41B represses T cell responsiveness in part via CD5

In analyzing scRNA-seq data, we observed that *Cd5*, a negative regulator of TCR signaling^45^, was downregulated in TMEM41B-deficient naive T cells (Figure 6A), a finding confirmed at the protein level through flow cytometry (Figure 6B). As CD5 represses TCR signaling, we hypothesized that TMEM41B-deficient T cells might be hyperreactive to antigen stimulation. Indeed, the upregulation of CD44, CD69 and CD25 was more pronounced in TMEM41B-deficient T cells compared to control T cells, especially at low doses of CD3 cross-linking (Figures 6C and S7A). Conversely, upregulation of surface proteins more dependent on NFAT signaling, including PD-1, TIM-3 and LAG-3, was diminished in TMEM41B-deficient T cells (Figures 6D and S7B), consistent with our initial screening data (Figures 1C, S1B and S1C).

**Figure 6.**
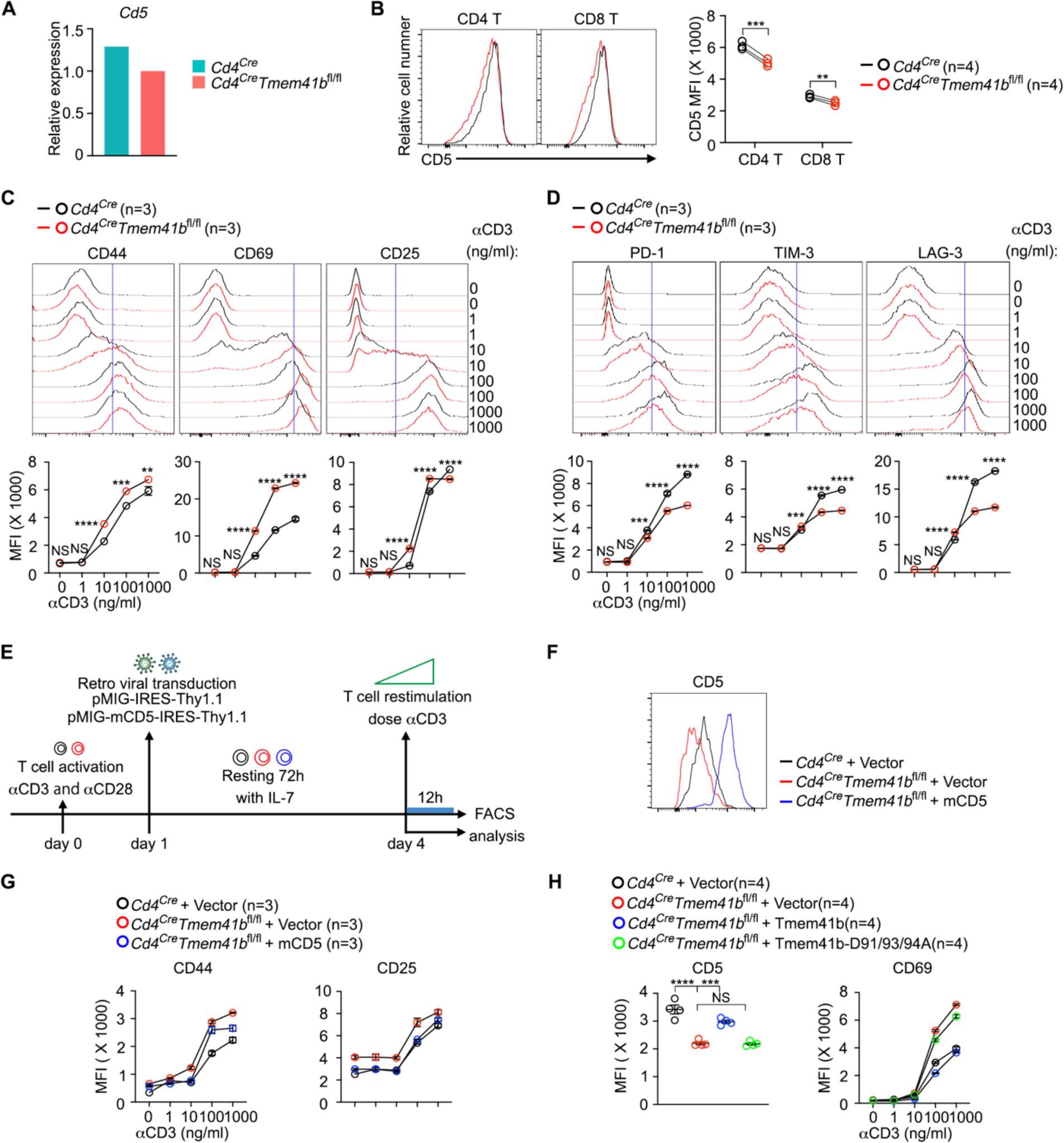
TMEM41B represses T cell responsiveness via CD5. (A) Relative mRNA levels of CD5 in control (*Cd4*^Cre^) and TMEM41B-deficient (*Cd4*^Cre^*Tmem41b*^fl/fl^) naive T cells (from sc-RNA seq data). (B) Flow cytometry analysis of CD5 expression on naive T cells in peripheral lymph nodes (pLN) of control and TMEM41B-deficient mice. Representative plots and statistics are shown. (n = 4 mice). (C) Flow cytometry analysis of the upregulation of CD44, CD69 and CD25 on CD4 T cells upon stimulation with different doses of αCD3 antibodies. Representative plots and statistics are shown. (n = 3 independent experiments). (D) Flow cytometry analysis of the upregulation of PD-1, TIM-3 and LAG-3 on CD4 T cells upon stimulation with different doses of αCD3 antibodies. Representative plots and statistics are shown. (n = 3 independent experiments). (E) Experimental design for CD5 rescue of phenotypes in TMEM41B-deficient T cells. (F) Flow cytometry analysis of CD5 expression on CD4 T cells transduced with indicated constructs. Representative plots are shown. (G) Flow cytometry analysis of CD44 and CD25 expression on CD4 T cells transduced with indicated constructs upon αCD3 stimulation. (n = 3 independent experiments). (H) Flow cytometry analysis of CD5 (before αCD3 stimulation) and CD69 (after αCD3 stimulation) on CD4 T cells transduced with indicated constructs. Statistics are shown. (n = 4 independent experiments). ** p< 0.01, *** p< 0.001, **** p< 0.0001, NS, not significant by two-tailed paired t-test (B), two-way ANOVA (C and D) or one-way ANOVA (H).

To explore whether reduced CD5 expression is the cause of hyperresponsiveness of TMEM41B-deficient T cells, we overexpressed CD5 in TMEM41B-deficient T cells (Figures 6E and 6F). Notably, the enhanced response of TMEM41B-deficient T cells to antigen stimulation was largely reversed by the overexpression of CD5 (Figures 6G and S7C), suggesting that CD5 downregulation contributes to the hyperresponsiveness of these cells.

To further examine the role of ER Ca^2+^ overload in CD5 downregulation and T cell hyperresponsiveness, we performed rescue experiments in TMEM41B-deficient T cells using the wild-type or D91/93/94A mutant of TMEM41B. Re-expression of the wild-type, but not D91/93/94A mutant of TMEM41B, largely reversed CD5 expression and hyperresponsiveness of TMEM41B-deficient T cells (Figure 6H), indicating that ER Ca^2+^ overload, through certain mechanism(s), downregulates CD5 in TMEM41B-deficient T cells, consequently heightening the responsiveness of these cells to antigen stimulation.

### TMEM41B-deficient T cells show reduced tolerance and heightened response to infections

Finally, we assess the consequences of TMEM41B deficiency in T cell responses in vivo. In a standard T cell tolerance test, wherein injection of αCD3 antibody induces T cell deletion (Figure 7A), TMEM41B-deficient T cells displayed relative resistance to αCD3-induced T cell deletion (Figures 7B and 7C), suggesting diminished tolerance of these cells. However, given that *Cd4^Cre^Tmem41b*^fl/fl^ mice did not manifest obvious signs of autoimmunity upon 6 months of age (data not shown), long-term monitoring is required to unveil potential spontaneous phenotypes.

**Figure 7.**
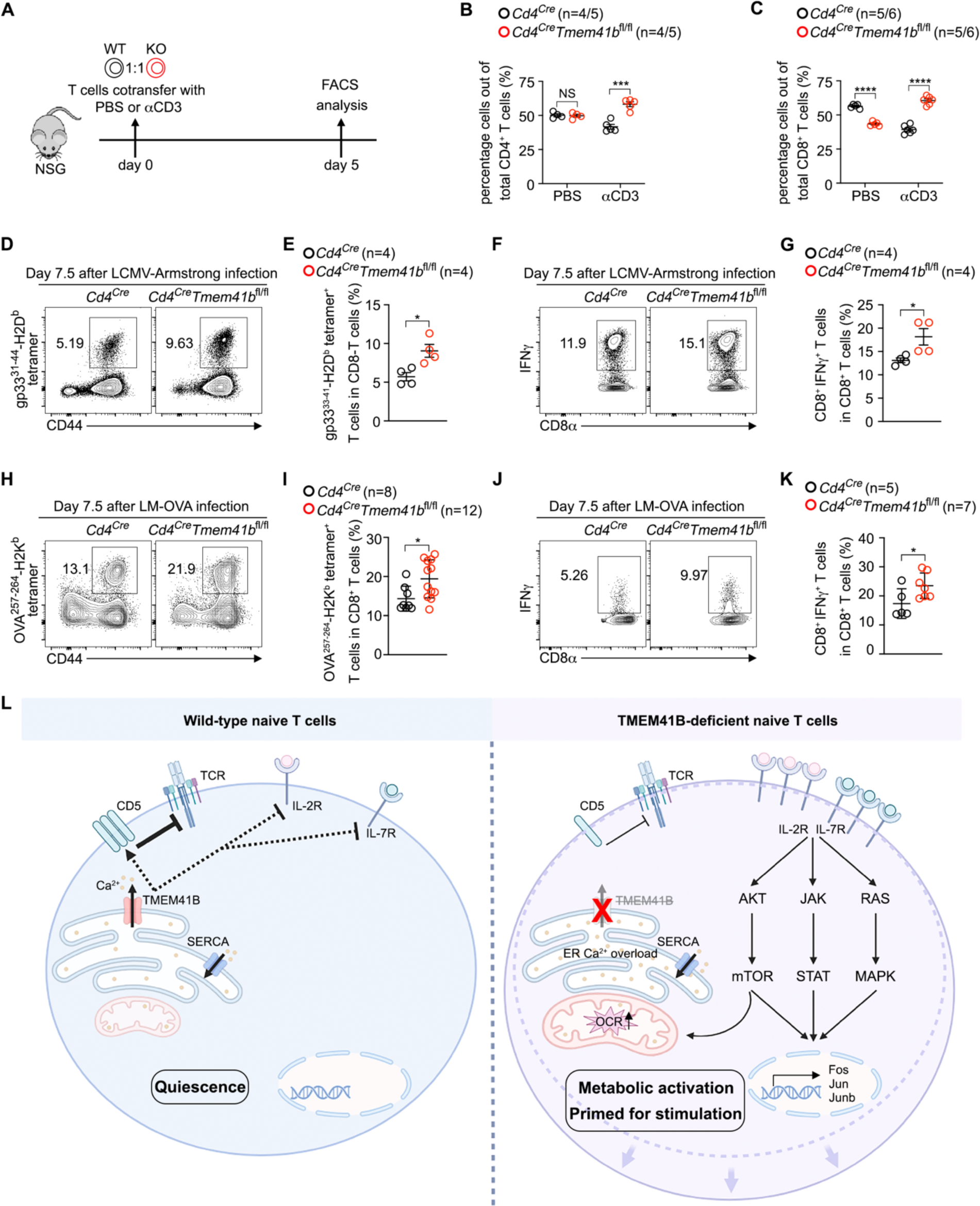
TMEM41B-deficient T cells exhibit reduced tolerance and increased response to infections. (A) Experimental design for in vivo T cell deletion by αCD3 antibodies. (B) Percentages of control (*Cd4*^Cre^) and TMEM41B-deficient (*Cd4*^Cre^*Tmem41b*^fl/fl^) CD4 T cells in NSG mice treated with PBS or αCD3 antibodies. (n = 4 or 5 mice). (C) Percentages of control and TMEM41B-deficient CD8 T cells in NSG mice treated with PBS or αCD3 antibodies. (n = 5 or 6 mice). (D and E) Flow cytometry analysis of H2D^b^-gp33^+^ CD8 T cells in spleens of control and TMEM41B-deficient mice 7.5 days after LCMV-Armstrong infection. Representative plots (D) and statistics (E) are shown. (n = 4 mice). (F and G) Flow cytometry analysis of IFNγ^+^ CD8 T cells (after ex vivo stimulation with gp33 peptide) in spleens of control and TMEM41B-deficient mice 7.5 days after LCMV-Armstrong infection. Representative plots (F) and statistics (G) are shown. (n = 4 mice). (H and I) Flow cytometry analysis of H2K^b^-OVA^+^ CD8 T cells in spleens of control and TMEM41B-deficient mice 7.5 days after Listeria monocytogenes (LM-OVA) infection. Representative plots (H) and statistics (I) are shown. (n = 8 or 12 mice). (J and K) Flow cytometry analysis of IFNγ^+^ CD8 T cells (after ex vivo stimulation with OVA^257-264^ peptide) in spleens of control and TMEM41B-deficient mice 7.5 days after LM-OVA infection. Representative plots (J) and statistics (K) are shown. (n = 5 or 7 mice). (L) A working model of TMEM41B Ca^2+^ channel in maintaining ER Ca^2+^ homeostasis and metabolic quiescence of naive T cells. * p< 0.05, *** p< 0.001, **** p< 0.0001, NS, not significant by two-tailed unpaired t-test (B, C, E, G, I and K).

We then explored the impact of TMEM41B on T cell response to infection. Although there was a slight reduction of naive CD8 T cells in *Cd4^Cre^Tmem41b*^fl/fl^ mice (Figure S3G), the percentages of antigen-specific CD8 T cells recognizing LCMV virus were significantly elevated in *Cd4^Cre^Tmem41b*^fl/fl^ mice compared to control mice during the peak response of LCMV Armstrong infection (Figures 7D and 7E). Upon ex vivo restimulation with the gp33 peptide, there were more IFNγ^+^ cells in *Cd4^Cre^Tmem41b*^fl/fl^ mice than control mice (Figures 7F and 7G). Similar observations were also made in a bacterial infection model with Listeria monocytogenes (LM)-OVA (Figures 7H - 7K). Thus, TMEM41B-deficient CD8 T cells displayed more robust responses in acute infections than control CD8 T cells, consistent with our in vitro findings that TMEM41B-deficient naive T cells are metabolically active and hyperreactive to antigen stimulation.

## DUSCUSSION

In this study, we demonstrate that TMEM41B functions as a concentration-dependent Ca^2+^ channel, releasing ER Ca^2+^ to prevent Ca^2+^ overload within the ER. ER Ca^2+^ overload drives TMEM41B-deficient T cells into a metabolically activated yet immunologically naive state, revealing a previously unappreciated but pivotal role of ER Ca^2+^ in maintaining metabolic quiescence and responsiveness of naive T cells, as depicted in Figure 7L.

The SERCA family ATPases continuously pump Ca^2+^ from cytosol into ER lumen^17^, establishing a substantial electrochemical gradient of Ca^2+^ between the ER and cytosol. In activated T cells and various other stimulated cell types, IP3R-mediated rapid release of ER Ca^2+^ store triggers SOCE^18^, a process crucial for T cell activation and metabolic reprogramming^19,20^. In naive T cells and other resting/quiescent cells, ER also passively releases Ca^2+21–23^, yet Ca^2+^ channel(s) responsible for such steady-state ER Ca^2+^ release was elusive^24–26^. Our data establish that TMEM41B is a bona fide ER Ca^2+^ release channel. Supporting evidences include: 1) TMEM41B deficiency resulting in ER Ca^2+^ overload (Figure 1); 2) overexpression of TMEM41B depleting ER Ca^2+^ (Figure 2); 3) overexpression of a plasma membrane-targeted TMEM41B inducing Ca^2+^ influx (Figure 2); 4) a Ca^2+^ channel activity of purified recombinant TMEM41B in an in vitro single-channel assay (Figure 3); 5) a TMEM41B mutant exhibiting altered Ca^2+^ channel activity (Figure 3). Thus, TMEM41B is a long-sought ER Ca^2+^ release channel that contributes to the passive release of ER Ca^2+^ under steady-state.

Given the vital roles of Ca^2+^ in physiology and the distinct concentrations of Ca^2+^ in various compartments, known Ca^2+^ channels are rigorously gated by diverse mechanisms, encompassing ligand binding, voltage sensitivity, mechanical force, and more. Our electrophysiologic data demonstrate that the opening of TMEM41B depends on Ca^2+^ concentration, exhibiting increased opening at higher Ca^2+^ concentrations. This characteristic aligns with the physiological role of TMEM41B in preventing ER Ca^2+^ overload. As ER Ca^2+^ levels surpass a certain threshold, deviating from the established setpoint or homeostasis, TMEM41B is activated to release ER Ca^2+^, thus averting the accumulation of harmful Ca^2+^ levels in the ER. Future structural studies will provide details of TMEM41B-meidated Ca^2+^ transport and the exact gating mechanism.

Biochemically, TMEM41B exhibits phospholipid scramblases activity^32,33^. Intriguingly, it is known that a single protein can serve dual functions as both an ion channel and a scramblase. For instance, the TMEM16 family of transmembrane proteins functions as both Ca^2+^-activated ion channels and phospholipid scramblases^46^. We speculate that TMEM41B, and potentially its counterparts TMEM41A, TMEM64 and VMP1 (TMEM49)^47^, may constitute another family of transmembrane proteins endowed with both ion channel and scramblase activities, warranting further researchs. In alignment with this notion, TMEM16 proteins have been shown to regulate SARS-CoV-2 infection by modulating Ca^2+^ oscillations in infected cells^48^. Notably, both TMEM41B and VMP1 are essential for the infection of various virus, including SARS-CoV-2^29–31^. It has been recently shown that Ca^2+^ microdomains on ER is essential for membrane budding^49^. Considering TMEM41B’s ability to release ER Ca^2+^ to cytosol and the reliance of viruses on host ER for vesicle budding, TMEM41B may control viral infection through its Ca^2+^ channel activity, which warrants future investigations.

Our previous research demonstrated the involvement of VMP1 in ER Ca^2+^ release in T cells^37^. Whether VMP1 is a Ca^2+^ channel remains to be determined. At cellular level, both TMEM41B- and VMP1-deficient cells manifest ER Ca^2+^ overload, and the overexpression of either protein depletes ER Ca^2+^, implying functional similarities between these two proteins. A noteworthy distinction is evident in VMP1-deficient T cells, where mitochondrial Ca^2+^ overload results in massive peripheral T cell death^37^. In contrast, TMEM41B-deficient T cells do not exhibit mitochondrial Ca^2+^ overload. It is possible that VMP1-deficient T cells experience more severe Ca^2+^ overload than TMEM41B-deficient cells, leading to the overflow of Ca^2+^ from the ER to mitochondria. Alternatively, VMP1 may localize at the ER-mitochondria junction, modulating ER-mitochondria Ca^2+^ transfer. Consequently, despite both TMEM41B and VMP1 contributing to ER Ca^2+^ release, the distinct consequences of ER Ca^2+^ overload in TMEM41B- and VMP1-deficient T cells may hinge on the extent of Ca^2+^ overload in the ER or the manner in which ER Ca^2+^ is released, which warrants future investigations.

The metabolic phenotype exhibited by TMEM41B-deficient T cells is unique. Unlike previously reported regulators of T cell quiescence, such as TSC1^50^, PTEN^51^ and Foxo1^52^, where metabolic activation was consistently accompanied by spontaneous T cell activation, TMEM41B-deficient T cells maintained an immunologically naive state while displaying significantly heightened metabolic activities. Marked increases in mitochondrial mass, OCR, ECAR, ROS levels, and cell size in TMEM41B-deficient naive T cells collectively point to metabolic activation, decoupled from immunological activation in these cells. Despite their immunological naivety, TMEM41B-deficient T cells exhibited features of “T cell activation”, including the downregulation of Klf2 and upregulation of Fos, Jun and Myc, suggesting these cells are in a transcriptionally primed state. Coupled with the downregulation of CD5, TMEM41B-deficient T cells are poised for antigen stimulation at both metabolic and signaling levels, explaining their heightened responsiveness to antigen stimulation, particularly at suboptimal antigen doses.

An important finding in this study is that the level of ER Ca^2+^ regulates the expression of key receptors on naive T cells, including IL-2 receptor α chain, β chain, common γ chain, IL-7Rα chain, and CD5. Despite a reduction in CD5 mRNA levels, no concurrent increase in mRNAs for IL-2 and IL-7 receptors was observed, suggesting that ER Ca^2+^ overload modulates the expression of the latter receptors at post-transcriptional level. The detailed mechanisms of such regulation await further investigation. Nevertheless, the increased expression of IL-2/IL-7 receptors and associated signaling events in TMEM41B-deficient T cells can only be reversed by wild-type TMEM41B but not a Ca^2+^ channel defective mutant (D91/93/94A), demonstrating that the metabolic activation of naive TMEM41B-deficient T cells is attributed to the Ca^2+^ channel activity and not other functions of TMEM41B. Thus, ER Ca^2+^ plays a previously unappreciated role in maintaining the metabolic quiescence of resting naive T cells by suppressing aberrant signaling from IL-2 and IL-7. Conversely, the increased responsiveness of TMEM41B-deficient T cells implies that targeting TMEM41B or ER Ca^2+^ represents a novel strategy to amplify T cell response during infections.

TMEM41B exhibits broad expression across various tissues. Although not specifically investigated in this study, it is plausible that TMEM41B-mediated ER Ca^2+^ release also plays a role in regulating the homeostasis of other immune cells. Beyond its impact on immune system, dysregulations in ER Ca^2+^ are implicated in a myriad of human diseases ^53,54^. The loss of TMEM41B causes spinal muscular atrophy (a neurodegenerative disease) in worms and mice^55,56^. Additionally, deletion of TMEM41B in the liver induces nonalcoholic hepatosteatosis in mice^32^. In humans, single nucleotide polymorphisms in TMEM41B are associated with viral infection such as SARS-CoV-2^29^. Targeting TMEM41B-mediated ER Ca^2+^ regulation holds promise for therapeutic interventions in these diverse pathological conditions.

## LIMITATIONS

Our study has unveiled TMEM41B as an ER Ca^2+^ release channel, whose activity is governed by Ca^2+^ concentration rather than voltage. To elucidate the precise Ca^2+^ transport and gating mechanism of this novel channel, follow-up studies involving detailed structural analysis are imperative. Further investigations are warranted to understand how ER Ca^2+^ homeostasis modulates IL-2/IL-7 receptors and CD5 in T cells. While we attribute the observed phenotypes of TMEM41B-deficient cells to ER Ca^2+^ overload, it remains plausible that the phenotypes are a result of the Ca^2+^ released from the ER by TMEM41B, either as microdomains on the ER surface or elsewhere, rather than ER Ca^2+^ per se. However, testing this hypothesis poses challenges due to technical limitations in selective compensating for Ca^2+^ released by TMEM41B without affecting ER Ca^2+^. Finally, although phenotypes of TMEM41B-deficient T cells can only be reversed by wild-type TMEM41B but not a Ca^2+^ channel activity-defective mutant, we cannot entirely exclude minor contributions of TMEM41B’s other functions to the observed T cell phenotypes. Alternatively, TMEM41B’s other functions may be downstream and/or dependent on its Ca^2+^ channel activity. The interplay and potential interconnections among TMEM41B’s three major functions: ER Ca^2+^ channel, autophagy, and lipid scrambling, merit thorough investigation in future studies.

## STAR★METHODS

### KEY RESOURCES TABLE

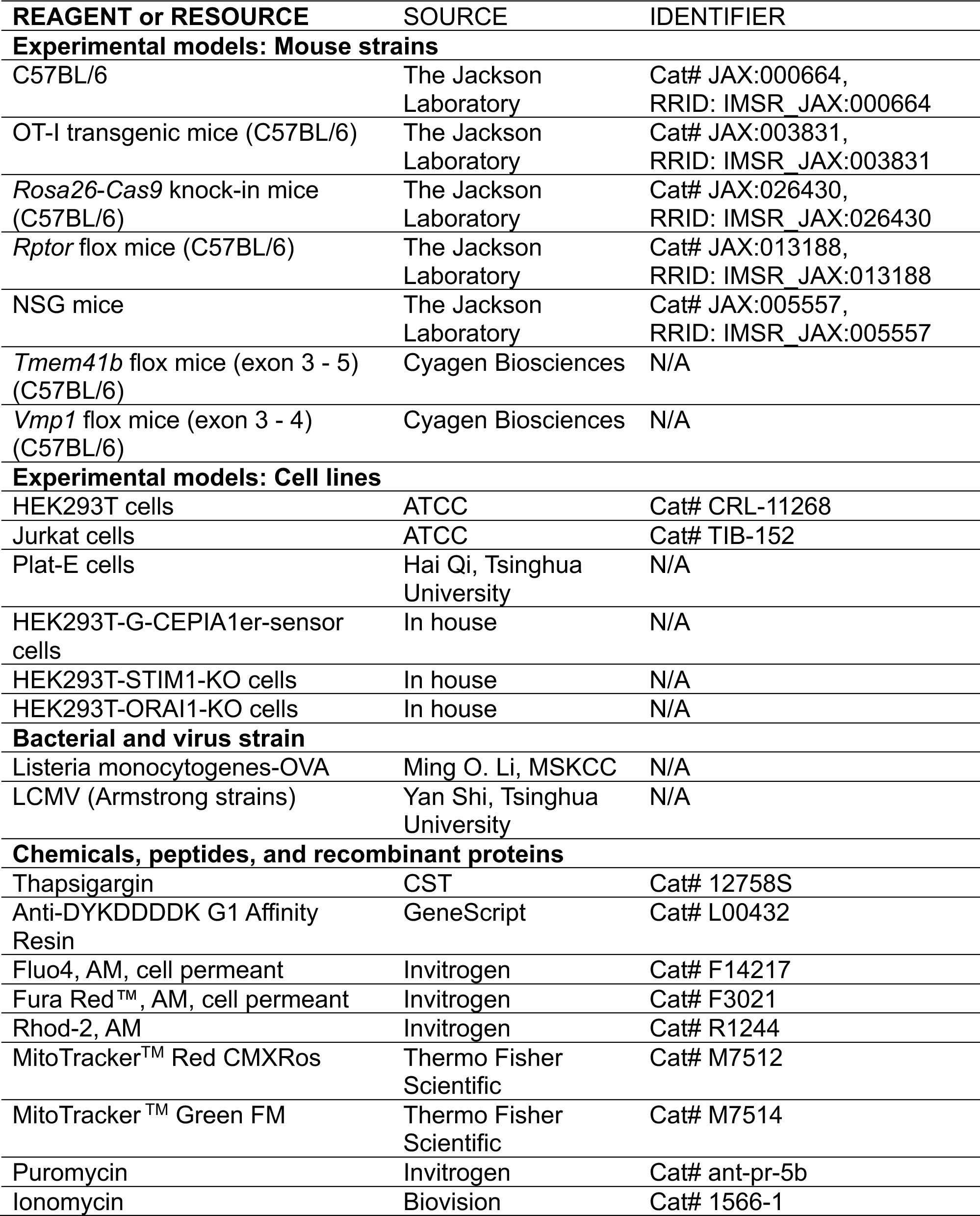
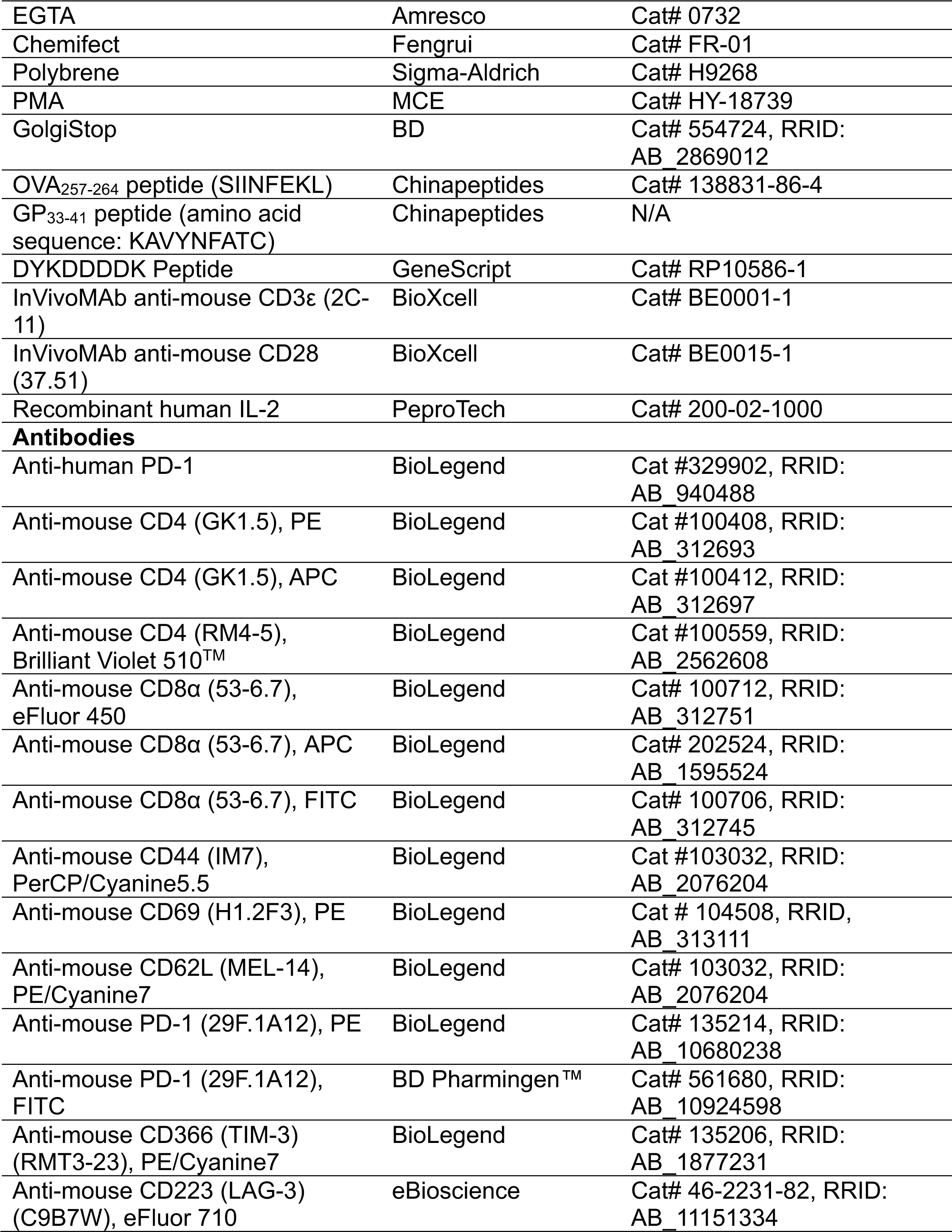
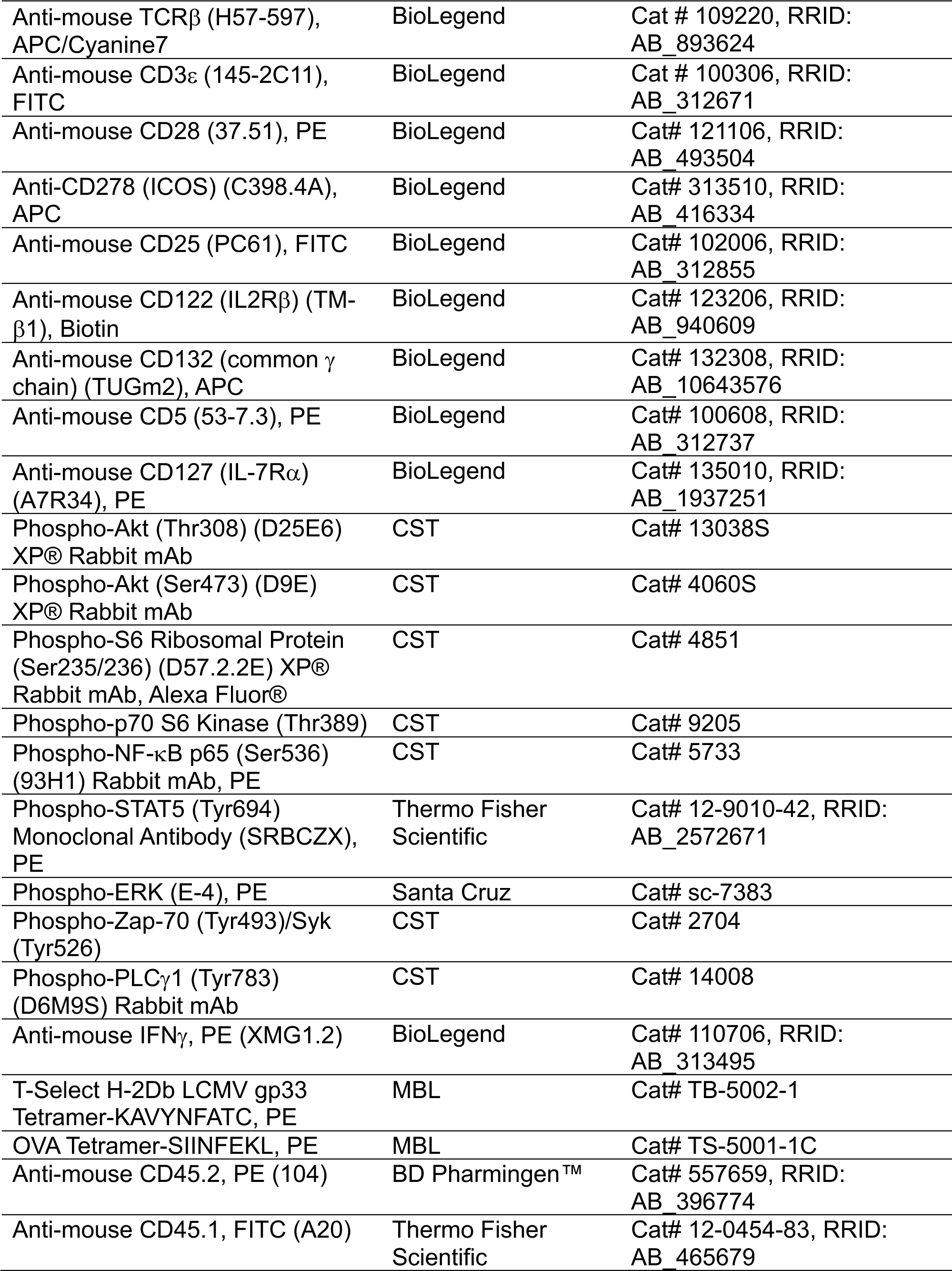
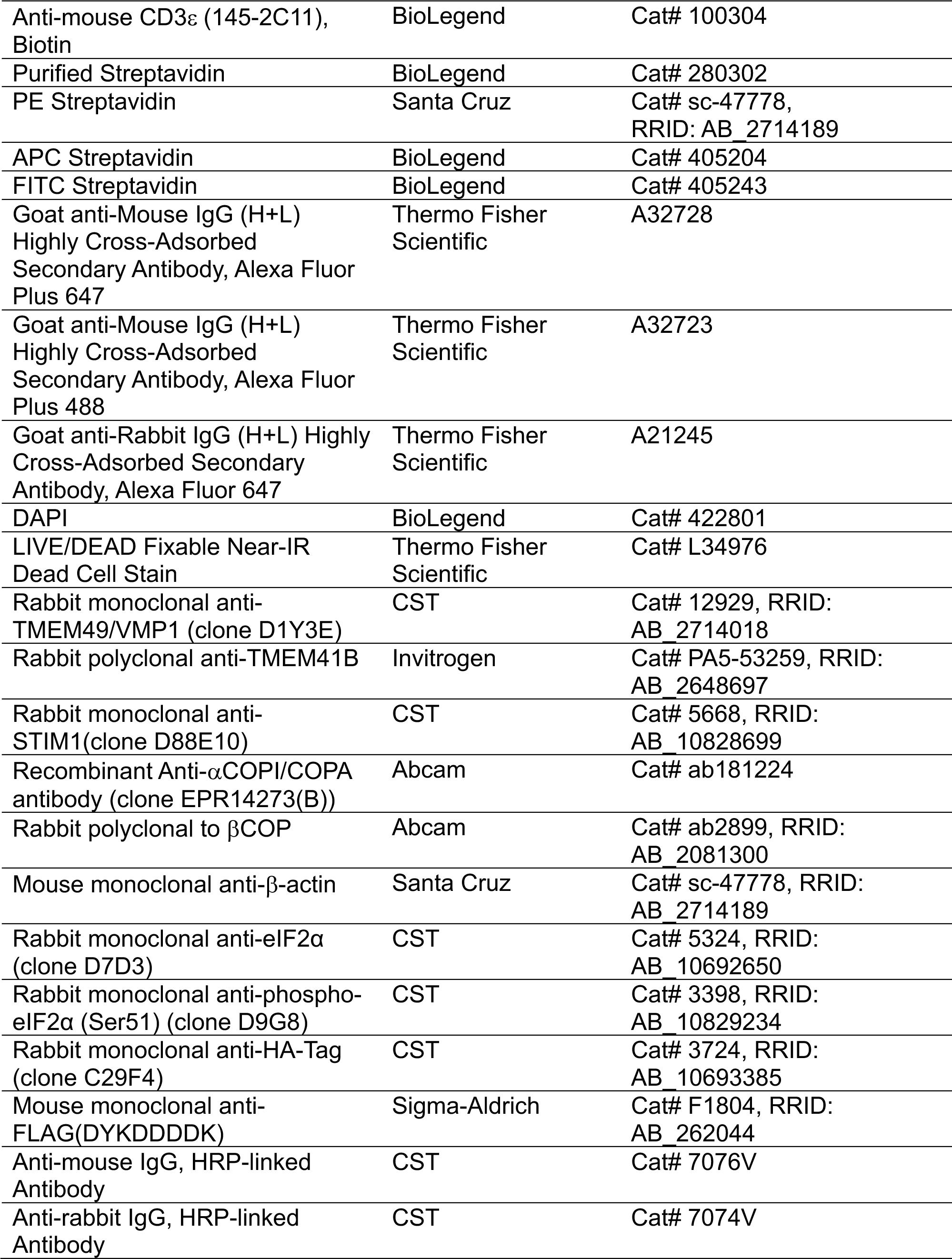
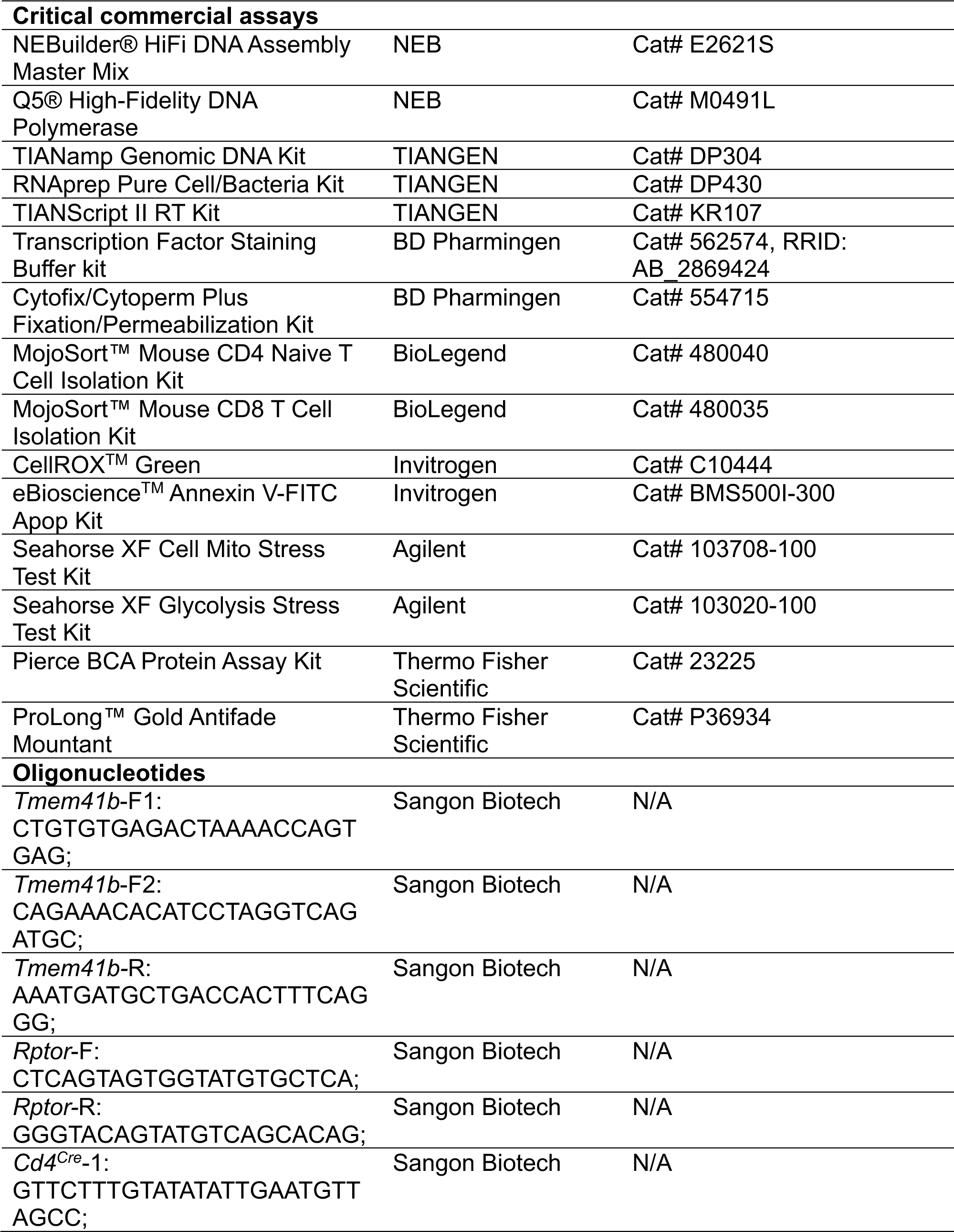
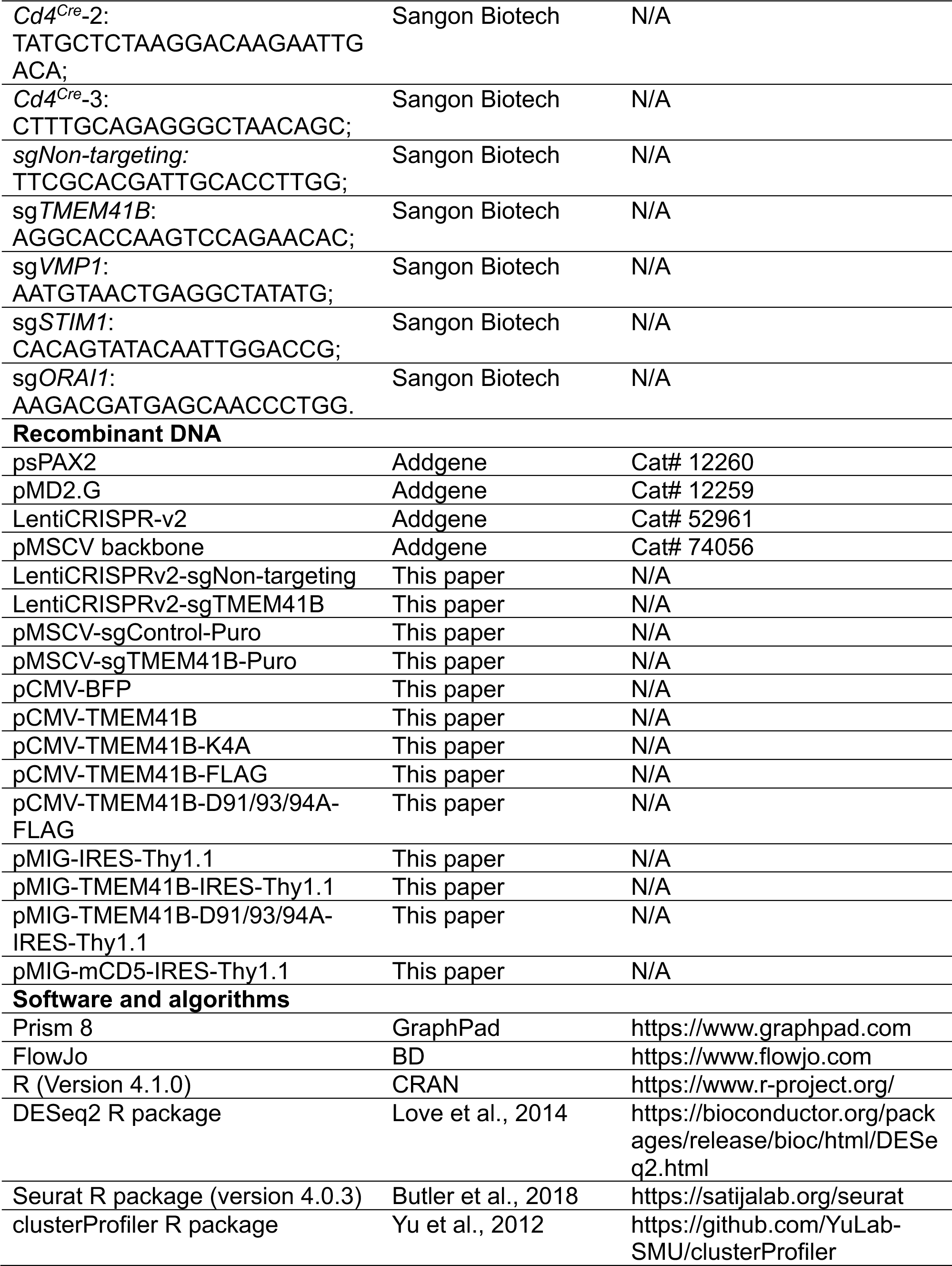
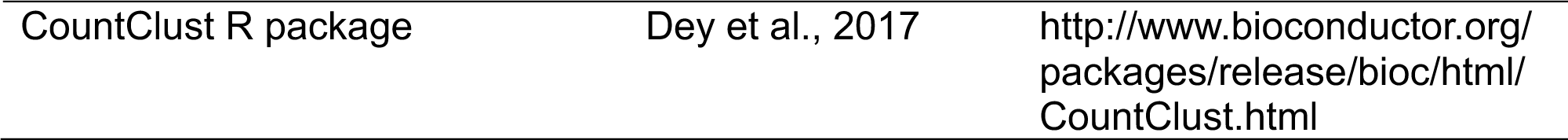

## RESOURCE AVAILABILITY

### Lead contact

Further information and requests for resources and reagents should be directed to and will be fulfilled by the lead contact, Min Peng.

### Materials availability

Animal strains used in this study are available from The Jackson Laboratory and Cyagen Biosciences.

### Data and code availability

- All data and code to understand and assess the conclusions of this research are available in the main text and supplementary materials. RNA-seq data will be deposited and publicly available.
- This paper does not contain original code.
- Any additional information required to reanalyze the data reported in this paper is available from the lead contact upon request.

## EXPERIMENTAL MODEL AND STUDY PARTICIPANT DETAILS

### Aminals

Cas9 mice (Stock No: 026430), OT-1 mice (Stock No: 003831), NSG mice (Stock No: 005557) and *Rptor* flox mice (Stock No: 013188) were from Jackson Laboratory. *Tmem41b* flox mice (exon 3 - 5) were generated by gene targeting service provided by Cyagen Biosciences. Genotyping primers were: *Tmem41b*-F: CTGTGTGAGACTAAAACCAGTGAG; *Tmem41b-*R: CAGAAACACATCCT AGGTCAGATGC; *Rptor*-F: CTCAGTAGTGGTATGTGCTCAG; *Rptor-*R: GGGTACAGTATGTCAGCACAG. *Cd4^Cre^* mice, *Tmem41b* flox mice and *Rptor* flox mice were C57BL/6J background. *Vmp1* flox mice have been reported previously^37^. Age and sex-matched littermates were used as control in all experiments. Mice were housed under specific pathogen-free conditions at the Laboratory Animal Research Center of Tsinghua University (Beijing, China). The facility was approved by Beijing Administration Office of Laboratory Animal. All animal works were approved by Institutional Animal Care and Use Committee (IACUC).

### Cell lines

HEK293T cells (CRL-11268) and Jurkat cells (TIB-152) were from ATCC. HEK293T cells were maintained in DMEM (Gibco) supplemented with 10% fetal bovine serum (FBS) (Gemini), 2 mM glutamine,100 units/ml of penicillin and 100 μg/ml of streptomycin at 37 °C in a humidified incubator with 5% CO_2_. Jurkat cells were maintained in RPMI 1640 (Gibco) with the same supplements mentioned above. All cell lines were tested for mycoplasma by the TransDect^TM^ PCR Mycoplasma detection Kit (TRAN, FM311), and were confirmed to be negative.

T cells were cultured in T cell medium (TCM): RPMI1640 medium (Gibco) supplemented with 5% FBS, 2 mM glutamine, 55 μM β-mercaptoethanol, 1 mM sodium pyruvate, 100 units/ml penicillin, 100 μg/ml streptomycin and 2 ng/ml IL-2. Cas9-OT-1 cells were activated by 1 μM OVA peptide (257-264, SIINFEKL) for 24 h, and passaged every 1 – 2 days at the density of 1 - 2 million cells/ml with 2 ng/ml IL-2. Primary T cells from spleen and lymph nodes were activated by 1 μg/ml anti-CD3 (BioXCell, BP0001-1, RRID: AB 1107634) and 1 μg/ml anti-CD28 (Bio X Cell, BE0015-1, RRID: AB_1107624) overnight, and passaged every 1 - 2 days at the density of 1 - 2 million cells/ml with 2 ng/ml IL-2.

## METHOD DETAILS

### Retrovirus production and viral transduction

Retrovirus were packaged by transfection of plat-E cells with associated plasmids using Chemifect^TM^ according to the manufacturer’s protocol. The viral supernatant was collected at 48- and 72-hours post-transfection, filtered via 0.45 μM filters, aliquoted and frozen at -80°C. Primary T cells were activated with 1 μg/ml αCD3 and αCD28, suspended in 1 ml TCM in 12-well plate. 2 ml Retrovirus and 8 μg/ml polybrene were added into T cells 20 ∼ 24 h post activation, followed by 2,000g centrifugation at 33 °C for 2 h, then the plate was returned to incubator. After 6 h of incubation, cells were transferred to 10-cm dish with fresh medium containing 2 ng/ml IL-2.

### CRISPR knockout of individual gene in Jurkat and HEK293T cells

Single guide RNAs were cloned into LentiCRISPRv2 (Addgene, #52961) to knockout specific gene in HEK293T and Jurkat cells. To generate polyclonal knockout cell lines, Lentivirus were produced in HEK293T cells by co-transfection of LentiCRISPRv2-sgRNA plasmids targeting specific gene together with psPAX2 and pMD2.G by Chemifect^TM^ according to the manufacturer’s protocol. For transduction, 3 million HEK293T or Jurkat cells were suspended in 1 ml fresh medium with 2 ml lentivirus and 8 μg/ml polybrene in 12-well plate, followed by 2,000g centrifugation at 33 °C for 2 h, then the plate was returned to incubator. After 6 h of incubation, cells were detached and transferred to 10-cm dish with fresh medium. Twenty-four hours after infection, puromycin (3 μg/ml) was added to cells for selection. Cells were selected for 3 days and expanded for another 3 days before experiments. The target sequences of sgRNA were: sgNon-targeting: TTCGCACGATTGCACCTTGG; sg*TMEM41B*: AGGCACCAAGTCCAGAACAC; sg*STIM1*: CACAGTATACAATTGGACCG; sg*ORAI1*: AAGACGATGAGCAACCCTGG.

To generate monoclonal knockout cell lines, HEK293T cells were transiently transfected with LentiCRISPRv2 plasmids targeting specific gene and selected with 3 μg/ml puromycin for 72 h. Then, cells were plated in 96-well plates with approximately one cell per well in DMEM with 15% FBS. After 10 days, single clones were expanded and verified by immunoblotting.

### CRISPR knockout of individual gene in Cas9-OT-1 cells

Single guide RNAs were cloned into pMSCV-sgRNA-Puro to knockout specific gene in Cas9-OT-1 cells. Retrovirus production, Cas9-OT-1 cells activation and spin-infection were performed as described above. Twenty-four hours after infection, puromycin (3 μg/ml) was added to cells for selection. Cells were selected for 3 days and expanded for another 2 days before experiments. The target sequence of sgRNA was: sg*Tmem41b*: GGCAGCAAAGATCATCTGAA.

### Ca^2+^ measurement with Fluo4/Fura Red by flow cytometry

Fura Red (Invitrogen, F3021) and Fluo4 (Invitrogen, F14217) were used for Ca^2+^ measurement according to published protocol^37^. In all Ca^2+^ assays, cells were plated at same density in 6-well plates one day before experiment to exclude the influence of cell density. One million control or gene-modified HEK293T, Jurkat or Cas9-OT-1 T cells resuspend in 250 μl Ca^2+^ assay medium (DMEM for HEK293T, RPMI 1640 for Jurkat and OT-1 T cells, and both medium were supplemented with 25 mM Hepes [pH = 7.4] and 2.5% FBS). Fura Red and Fluo4 were mixed together and prepared as 2 × solution in 250 μl Ca^2+^ assay medium. Then, the 250 μl medium with cells and 250 μl medium with 2 × Fura Red and Fluo4 mixture were mixed together, and the final concentration for Fura Red and Fluo4 were both 2 μM. The mixtures were incubated at 37 °C for 30 min in the dark. Cells were washed once with Ca^2+^ assay medium and resuspend in 800 μl Ca^2+^ assay medium for flow cytometry analysis.

For measurement of thapsigargin-induced Ca^2+^ influx directly, we placed cell suspension on the cytometer, adjusted the baseline intensities for Fluo4 (FITC channel) and Fura Red (PerCP-Cy5.5 channel), and recorded for 60 s. Then we removed the tube from the cytometer (without stopping the acquisition/recording), quickly added thapsigargin to a final concentration of 10 nM or 1 μM as indicated. Samples were quickly mixed, returned to the cytometer. The total assay time was indicated in each experiment.

For measurement of ER Ca^2+^ and SOCE, cell suspension was supplemented with 2 mM final concentration of EGTA before placed on cytometer. The concentration of Ca^2+^ is 0.42 mM in RPMI 1640 and 1.8 mM in DMEM, so 2 mM EGTA can completely chelate the extracellular Ca^2+^. We adjusted the baseline intensities for Fluo4 (FITC channel) and Fura Red (PerCP-Cy5.5 channel), and recorded for 60 s. Then we removed the tube from the cytometer, quickly added thapsigargin to a final concentration of 1 μM. Samples were quickly mixed, returned to the cytometer and recorded for another 420 s. Then, we removed the tube from the cytometer (without stopping the acquisition/recording), quickly added CaCl_2_ to a final concentration of 2 mM (The Ca^2+^ in assay medium was not included in this 2 mM concentration). Samples were quickly mixed, returned to the cytometer and recorded for another 420 s. All measurements were performed at 25°C.

For measurement of Ca^2+^ influx in cells transfected with plasma membrane-targeted TMEM41B, dye-loaded cells were recorded for 60 s. Then we removed the tube from the cytometer (without stopping the acquisition/recording), quickly added 8 mM or indicated concentration of CaCl_2_. Samples were quickly mixed, returned to the cytometer and recorded for another 240 s.

For measurement of ER Ca^2+^ store in WT and TMEM41B-deficient T cells, freshly isolated T cells were loaded with Fluo4 (FITC channel) and Fura Red (PerCP-Cy5.5 channel). Then cell suspension was supplemented with 2 mM final concentration of EGTA before placed on cytometer and recorded for 60 s. Then we removed the tube from the cytometer (without stopping the acquisition/recording), quickly added 1 μM Ca^2+^ ionophore ionomycin. Samples were quickly mixed, returned to the cytometer and recorded for another 240 s.

For measurement of TCR-induced SOCE in WT and TMEM41B-deficient T cells, freshly isolated T cells were incubated with 10 μg/ml αCD3-biotin, then 10 μg/ml purified streptavidin (STV) was used for cross-linking. SOCE was examined by flow cytometry as above described.

### Ca^2+^ measurement using Ca^2+^ sensors G-CEPIA1er by flow cytometry

Cell line stably expressing sensors were established by lentiviral transduction^37^. TMEM41B were knocked out in this cell line by methods described above. For measurement of the fluorescence of G-CEPIA1er, one million cells were suspended in 800 μl Ca^2+^ assay medium. We placed cell suspension on the cytometer and G-CEPIA1er (GFP channel) was recorded for 60 s. Then we removed the tube from cytometer, quickly added thapsigargin to a final concentration of 1 μM. Samples were quickly mixed, returned to the cytometer and recorded for another 840 s. The total assay time for each sample was 900 s. The voltages for GFP (for G-CEPIA1er) channels were kept unchanged for all samples. In overexpression studies, sensor cells were transfected with indicated constructs with a BFP reporter, and the influence of overexpression on the intensity of sensor was measured on BFP^high^ cells.

### Overexpression experiments

To overexpress proteins in HEK293T cells, the full-length cDNAs of human TMEM41B was cloned into a small vector pCMV with a backbone of 3.7 kb. This vector does not contain fluorescent marker, so a pCMV-BFP construct was co-transfected to gate out BFP-high cells for analysis on flow cytometry.

To overexpress proteins in T cells, the full-length cDNAs of mouse TMEM41B was cloned into pMIG vector with Thy1.1 as a reporter marker. Retrovirus production, T cells activation and spin-infection were performed as described above. 72h post spin infection, Thy1.1 positive T cells were gated out for analysis on flow cytometry.

### Immunofluorescence

HEK293T cells were plated on poly-d-lysine Cellware 12-mm coverslips (BD Biosciences). On the next day, cells were transfected with indicated plasmids. Twenty-four hours after transfection, cells were rinsed once with PBS, and fixed for 15 min with 4% para-formaldehyde in PBS at room temperature. Cells were rinsed twice with PBS and permeabilized by methanol (pre-chilled at -20°C) for 10 min on ice. After rinsing three times with PBS, the slides were blocked by 5% BSA for 30 min at room temperature, and incubated with primary antibodies (diluted 1:500 in 5% BSA) for 1 - 3 h at room temperature, then rinsed four times with PBS, and incubated with goat-anti-mouse Alexa Fluor secondary antibodies (diluted 1:1,000 in 5% BSA) for 1 h at room temperature in the dark. Slides were washed four times with PBS, mounted on glass coverslips using ProLong Gold (Invitrogen) with DAPI, and imaged on a Zeiss780 confocal microscope with a 63 × oil lens. The anti-FLAG antibody used for immunofluorescence staining was from Sigma (M2, F1804).

### Purification of recombinant TMEM41B proteins

HEK293T cells were plated in a 15-cm plate 16 h before transfection. Twenty μg of plasmids encoding FLAG-TMEM41B and FLAG-TMEM41B-D91/93/94A were transfected using Chemifect^TM^ according to the manufacturer’s protocol. After 48 h of transfection, cells were collected, washed twice with ice-cold PBS, and lysed with 5 packed cell volumes of 1% DDM lysis buffer (20 mM Tris-Cl, [pH 8.0], 2 mM CaCl_2_, 200mM NaCl, 1% DDM, 0.2% (w/v) CHS, 0.012% (w/v) glyco-diosgenin (GDN, Anatrace) and 1×EDTA-free protease inhibitor cocktail) for 2h with gentle rotation at 4°C. The lysate was spun at 15,000 rpm for 30 min and the supernatant was incubated with 100 μl (packed volume) of anti-DYKDDDDK G1 Affinity resin (GenScirpt) for 3 h with gentle rotation at 4°C. The beads were washed five times with wash buffer (20 mM Tris-Cl, [pH = 8.0], 200 mM NaCl, 2mM CaCl_2_, 0.02% GDN). Bound proteins were eluted with 500 μl wash buffer containing 0.125 mg ml^−1^ FLAG peptide (GenScirpt) for 2 h with gentle rotation at 4°C. Buffer containing eluted protein was filtered through a 0.22-μm spin filter and directly loaded on the Enrich SEC650 column (Bio-Rad) for SEC. Fractions containing the proteins were analyzed.

### Western blot

After treatment as indicated, cells were collected and washed with cold-PBS. Cells were lysed with lysis buffer (1% Triton X-100, 40 mM HEPES, [pH 7.4], 10 mM ϕ3-glycerol phosphate, 10 mM pyrophosphate) supplemented with EDTA-free protease inhibitor cocktail (Thermo-Fisher) on ice for 15 min. The soluble fractions of cell lysates were isolated by centrifugation at 15,000 rpm for 10 min at 4 °C. Proteins were denatured by the addition of 6 × SDS sampling buffer and boiling for 10 min at 95 °C. Samples were subjected to SDS-PAGE and immunoblotting analysis.

### Electrophysiology recording of TMEM41B by Planar Lipid Bilayers

The electrophysiological experiment was performed with the Planar Lipid Bilayer Work station (BLM Workstation). Chamber and Cup separate the solution into two compartments that were connected with a 200 μm hole. We define Cup as the *Cis* side, and Chamber as the *Trans* side. 500 mM and 50 mM KCl solutions were added in *Cis* and *Trans* side respectively. The phospholipids (phosphatidylcholine: phosphatidylserine= 3: 2) dissolved in decane were painting softly into the small hole in Cup to form lipid bilayer. All lipids were bought from Avanti (Avanti Polar Lipids, USA). Purified TMEM41B or mutant protein were added in *Cis* side to incorporate into planar lipid bilayer. Single channel currents were recorded under voltage-clamp mode using a Warnner bilayer clamp amplifier BC-535 (Warner Instruments, USA), filtered at 1 – 2 kHz.

The recording frequency was 10 kHz. Analog voltage was digitized with Digidata 1440A (Molecular Devices, US), and the data was stored by pCLAMP 10.4. The amplitude of Events and open probability (P_open_) were detected by Clampfit. Events with opening time less than 1.5 ms were ignored. Single-channel conductance was determined by fitting to Gaussian functions equations. The equilibrium potential was calculated using the Nernst equation and Goldman–Hodgkin–Katz flux equation.

### Flow cytometry

Singe cell suspensions were prepared from thymi, spleens, and lymph nodes by grinding through 70 μm strainer. Erythrocytes were depleted by hypotonic lysis. For staining of surface markers, cells were incubated in FACS buffer (PBS supplemented with 1% calf serum, 1% penicillin/streptomycin and 2 mM EDTA) with indicated combinations of antibodies for 15 min at 4°C together with Fc blockade (2G4) to prevent non-specific binding. Cells were washed twice with FACS buffer, DAPI was included to exclude dead cells.

For apoptosis assays, T cells were stained with surface marker, then Annexin V staining was performed following the instructions of Annexin V-FITC Apop Kit (Invitrogen: BMS500FI-300). Samples were recorded with an LSR Fortessa cytometer (BD) and analyzed with FlowJo software (BD).

For measurement of mitochondrial Ca^2+^, cells were washed with serum free RPMI 1640 medium and incubated with 1 μM Rhod-2-AM (Invitrogen, R1244) together with antibodies for CD4 (GK1.5) and CD8 (53-6.7) at 37 °C for 30 min. Cells were washed twice and resuspended in serum free RPMI medium, DAPI was used to exclude dead cells. Samples were acquired with an LSR Fortessa cytometer (BD).

For measurement of mitochondrial mass and mitochondrial membrane potential, cells were incubated with 200 nM MitoTracker® Probes (MitoTracker® Green for mitochondrial mass, MitoTracker® Red for mitochondrial membrane potential) together with antibodies for CD4 (GK1.5) and CD8 (53-6.7) at 37 °C for 30 min. After incubation, cells were washed twice and subjected to flowcytometry analysis. DAPI was used to exclude dead cells.

For measurement of total reactive oxygen spices (ROS), cells were incubated with total ROS probe (Invitrogen; Cat. # 88-5930) together with antibodies for CD4 (GK1.5) and CD8 (53-6.7) at 37 °C for 30 min. After incubation, cells were subject to flowcytometry analysis. DAPI was used to exclude dead cells.

For phospho-flow assays, T cells were first stained with surface markers togerher with LIVE/DEAD Fixable Near-IR (Cat# L34976, Invitrogen) at 4 °C for 15 min. Cells were washed twice with FACS buffer. Permeabilization was performed with permeabilization buffer from Transcription Factor Staining Buffer kit according to manufacturer’s instructions (BD Pharmingen). Then cells were stained with antibodies for 30 min at room temperature. After 2 washes, samples were analyzed by a LSR Fortessa cytometer (BD).

### Single-cell RNA sequencing and analysis

Naive T cells (TCRβ^+^CD62L^+^CD44^-^CD25^-^) were sorted by flow cytometry, and single cell suspensions were directly loaded on a microfluidic chip (Singleron GEXSCOPETM Single Cell RNA-seq Kit, Singleron Biotechnologies) and processed for scRNA-seq library preparation according to manufacturer’s protocol. The ultimate constructed and purified library with the target recovery of ∼ 13,000 single cells was sequenced on Illumina novaseq 6000. CeleScope (v1.9.0) pipeline was used for scRNA-seq data alignment and quantification. The generated data files including aligned and filtered reads, barcodes and unique molecular identifiers were processed by Seurat (v4.3.0) for further analysis. For the data set, cells were considered as low-quality and then excluded if number of detected genes < 200 or > 3,000. Cells were also removed if their mitochondrial gene proportions were larger than 10%. Following normalization process, the top 2,000 variable genes were chosen for principle-component analysis. 1 - 10 PCs, determined by *JackStraw* function as significant ones, were selected for UMAP and clustering analysis. Cluster specific genes were identified using *FindAllMarkers* (log FC threshold = 0.25) function. *FeaturePlot* and *VlnPlot* functions were also used for data visualization. For signature geneset scoring, *AddModuleScore* function from Seurat was used by selected gene sets.

### Seahorse experiments

Oxygen consumption rate (OCR) and extracellular acidification rate (ECAR) were measured with an XF96 extracellular flux analyzer (Seahorse Bioscience) according to the manufacturer’s instructions. Briefly, freshly isolated naive T cells were seeded on XF96 microplates (150,000 cells/well) that had been pre-coated with Cell-Tak adhesive according to the manufacturer’s instructions (BD Biosciences). The Mito stress test kit (Seahorse Biosciences) was used to test OCR under different conditions. Firstly, cells were incubated in the mito stress test medium without any drugs, and four baseline recordings were assessed. Then maximal OCR was obtained by sequential injection of 1 μM oligomycin and 0.25 μM FCCP that uncoupled oxygen consumption from ATP. Finally, 0.5 μM rotenone/antimycin A that inhibited complex I and III was injected. Glycolysis was measured with XF glycolysis stress test kit (Seahorse Biosciences). Initially, cells were incubated in the glycolysis stress test medium without glucose, and four basal ECAR recordings were assessed. Following by sequential injection of 10 mM glucose and 1 μM oligomycin that inhibited mitochondrial ATP production, the energy production shifted to glycolysis. The increased ECAR revealed the maximum glycolytic capacity of T cells. Finally, 100 mM 2-DG was injected to inhibited glycolysis.

### αCD3 induced T cell deletion in vivo

Naive CD4 or CD8 T cells were purified using FACS sorting. Wild-type and TMEM41B-deficient T cells were mixed at a 1:1 ratio (a total of 1 × 10^6^ cells), and co-transferred into NSG mice via tail vein. Mice were then injected with 10 mg of anti-CD3χ antibodies or PBS control. Donor T cells in spleen were analyzed by flow cytometry on day 5 post-transfer.

### LCMV Armstrong infection

Mice were infected with LCMV Armstrong virus (1×10^5^ PFU) by intra-peritoneal injection. On day 7.5 post-infection, mice were sacrificed and splenocytes were isolated to measure antigen-specific CD8 T cell response by flow cytometry. Briefly, splenocytes were stained with CD8α-PB, TCRβ-APC, CD44-Percp-Cy5.5 together with PE conjugated H-2D^b^-gp^33–41^-tetramer for 30 min at room temperature. Cells were washed twice with FACS buffer and analyzed by a LSR Fortessa cytometer (BD). Dead cells were excluded by DPAI staining.

### Listeria monocytogenes infection

5,000 colony-forming units (CFUs) of listeria monocytogenes expressing the chicken ovalbumin (LM-OVA) were injected to mice via tail vein. On day 7.5 post-infection, mice were sacrificed and splenocytes were isolated to measure antigen-specific CD8 T cell response by flow cytometry as described above.

## QUANTIFICATION AND STATISTICAL ANALYSIS

The investigators were not blinded to the treatment groups. Data are presented as means ± s.e.m. The P values and number of replicates (n) are shown in figures or figure legends. N represents the number of animals for in vivo/ex vivo experiments. For in vitro cell culture experiments, N indicates the number of independent experiments. GraphPad Prism 8.0 was used for statistical analysis. Paired or unpaired two-tailed Student’s t test, one-way ANOVA or two-way ANOVA were used to evaluate the difference between groups. P < 0.05 was considered significant. Representative flow plots, immunoblots, and micrographs were selected from biological replicates.

## SUPPLEMENTAL INFORMATION

This manuscript contains 7 supplemental figures.

## ACKNOWLEDGMENTS

We thank H. Qi’ lab, Y. Shi’s lab and B. Xiao’s lab for helps with calcium assays. We thank Institute for Immunology at Tsinghua University for providing and maintaining equipment.

This research was supported by Vanke Special Fund for Public Health and Health Discipline Development Tsinghua University (NO.2022Z82WKJ013, to M. P.), Tsinghua University DUSHI Program (52302102323, to M.P.), Tsinghua-Peking Center for Life Sciences (to M. P.), SXMU-Tsinghua Collaborative Innovation Center for Frontier Medicine (to M. P.), National Science Fund of Distinguished Young Scholars (81825021) (to Z.G.) and Shanghai Rising-Star Program (22QA1411000) (to B.X.).

## AUTHOR CONTRIBUTIONS

Y. M. performed experiments, analyzed data and prepared figures; Y. W. performed BLM experiments and prepared relevant figure (Figure 3 and related S3) under the guidance of B. X. and Z. G.; X. Z. contributed to the purification of recombinant TMEM41B; G. J. analyzed RNA-seq data and prepared relevant figures; J. X. and Z. L. contributed to experiments and management of mouse colony; N. Y. contributed to the design and supervision of the study; B. X. contributed to and supervised the BLM experiments; Z. G. supervised the BLM experiments and acquired funding; M. P. conceptualized and coordinated the study, acquired funding, analyzed the data and wrote the paper. All authors read and edited the manuscript.

---

**Figure S1.**
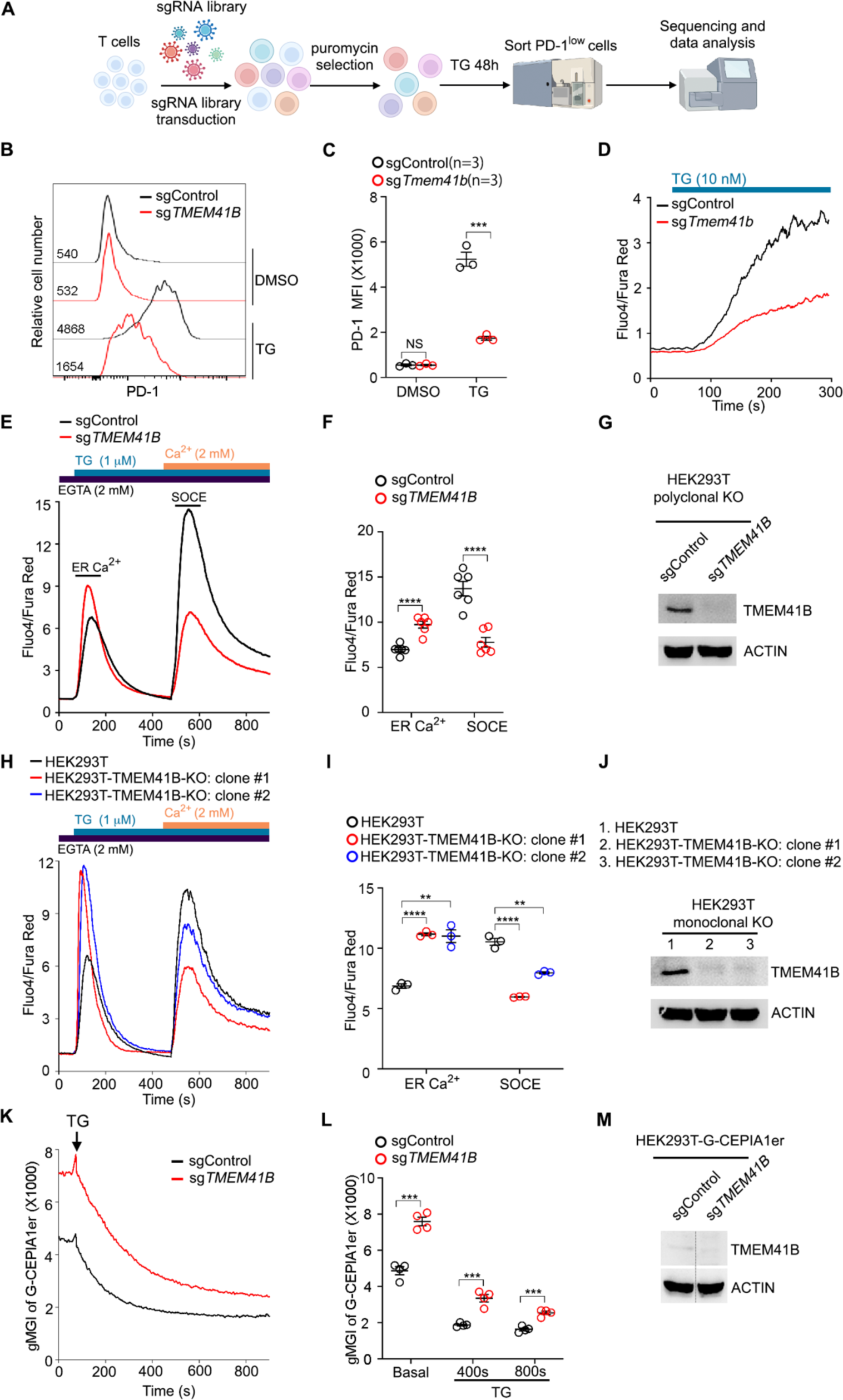
TMEM41B deficiency causes ER Ca^2+^ overload, related to Figure 1. (A) A flow chart of PD-1-based CRISPR screening of ER Ca^2+^ regulators. (B and C) Flow cytometry analysis of PD-1 expression on OT-1 cells (Cas9^+^) expressing indicated sgRNAs upon treatment with DMSO or thapsigargin (TG, 10 nM) for 48 h. Representative plots (B) and statistics (C) are shown. (n = 3 independent experiments). (D) Flow cytometry analysis of TG-induced Ca^2+^ influx in OT-1 cells (Cas9^+^) expressing indicated sgRNAs. Representative plots from 3 independent experiments are shown. (E and F) Flow cytometry analysis of ER Ca^2+^ and store-operated Ca^2+^ entry (SOCE) in HEK293T cells (Cas9^+^) expressing indicated sgRNAs. Representative plots (E) and statistics (F) are shown. (n = 6 independent experiments). (G) Immunoblot of TMEM41B in HEK293T cells (Cas9^+^) expressing indicated sgRNAs. ACTIN served as loading control. Representative blots from 2 independent experiments are shown. (H and I) Flow cytometry analysis of ER Ca^2+^ and SOCE in control and monoclonal TMEM41B knockout HEK293T cells. Representative plots (H) and statistics (I) are shown. (n = 3 independent experiments). (J) Immunoblot of TMEM41B in control and TMEM41B knockout HEK293T cells. ACTIN served as loading control. Representative blots from 2 independent experiments are shown. (K and L) Flow cytometry analysis of ER Ca^2+^ in HEK293T-G-CEPIA1er cells (Cas9^+^) expressing indicated sgRNAs. Representative plots (K) and statistics (L) are shown. (n = 4 independent experiments). (M) Immunoblot of TMEM41B in HEK293T-G-CEPIA1er cells (Cas9^+^) expressing indicated sgRNAs. ACTIN served as loading control. Representative blots from 2 independent experiments are shown. ** p< 0.01, *** p< 0.001, **** p< 0.0001, NS, not significant by two-tailed unpaired t-test (C, F and L) or one-way ANOVA (I).

**Figure S2.**
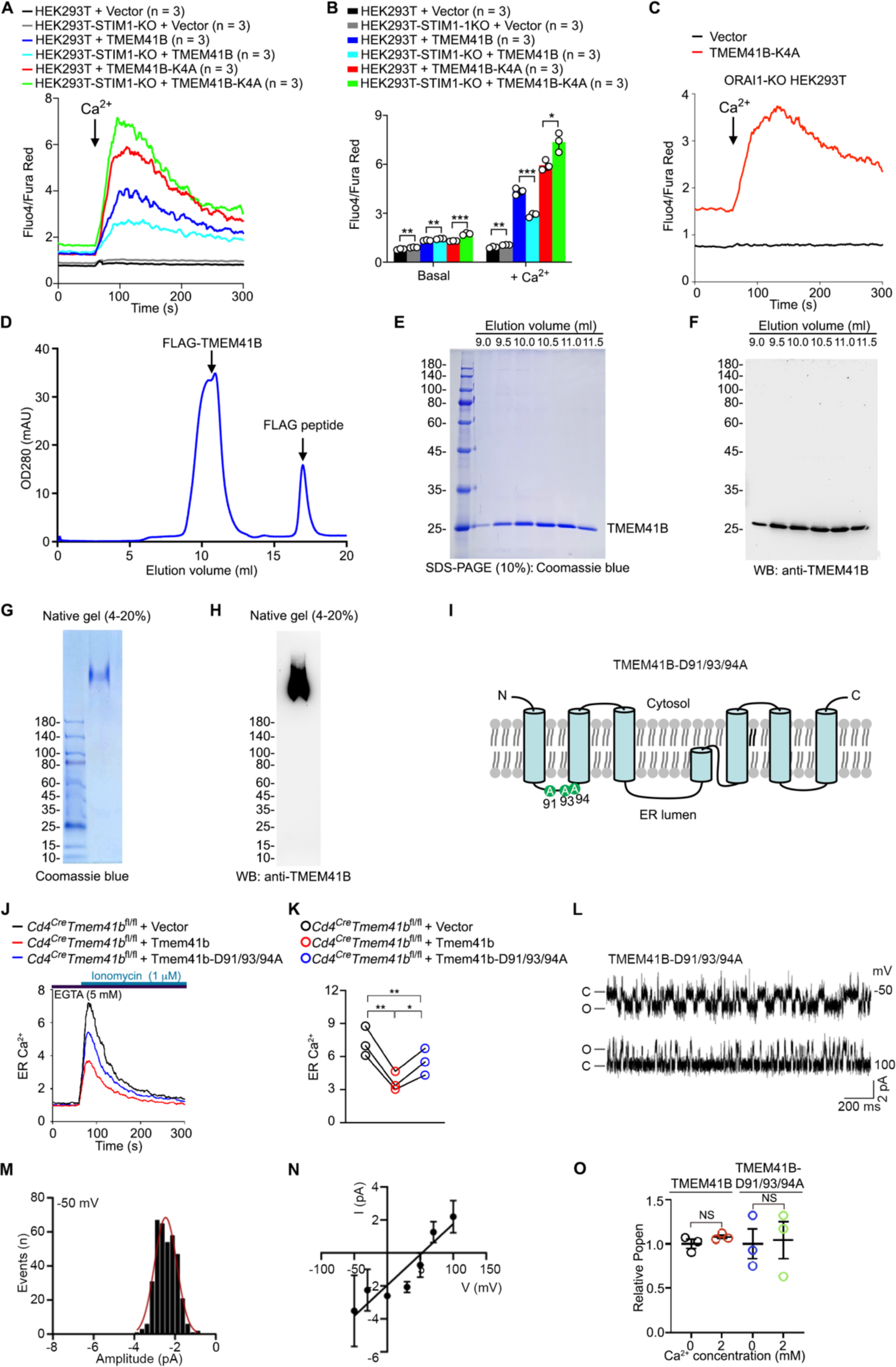
Biochemical and functional study of TMEM41B, related to Figure 2 and Figure 3. (A and B) Flow cytometry analysis of Ca^2+^ influx in control and STIM1-knockout (KO) HEK293T cells transfected with indicated plasmids. Representative plots (A) and statistics (B) are shown. (n = 3 independent experiments). (C) Flow cytometry analysis of Ca^2+^ influx in ORAI1-KO HEK293T cells transfected with indicated plasmids. Representative plots from 3 independent experiments are shown. (D) A representative plot of size exclusion chromatography (SEC) of purified TMEM41B. (E) Coomassie blue staining of TMEM41B from indicated elution volume of SEC. (F) Immunoblotting of TMEM41B from indicated elution volume of SEC. (G) Coomassie blue staining of TMEM41B separated by native gel. (H) Immunoblotting of TMEM41B separated by native gel. (I) Putative topology of TMEM41B protein. D91/93/94 mutations are shown. Mutated protein was named TMEM41B-D91/93/94A. (J and K) Flow cytometry analysis of ER Ca^2+^ in TMEM41B-deficient CD4 T cells retrovirally expressing indicated proteins. Representative plots (J) and statistics (K) are shown. (n = 3 independent experiments). (L) Representative single channel currents of TMEM41B-D91/93/94A in 500: 50 KCl solution at indicated voltage. (M) All-point current histograms for the trace in (L). (N) I-V curve of TMEM41B in solution of (L). (O) Relative open probability of TMEM41B and TMEM41B-D91/93/94A in (Figure 3K). Representative data from 3 independent experiments are shown (D - H). * p< 0.05, ** p< 0.01, *** p< 0.001, NS, not significant by two-tailed unpaired t-test (B and O) or one-way ANOVA (K).

**Figure S3.**
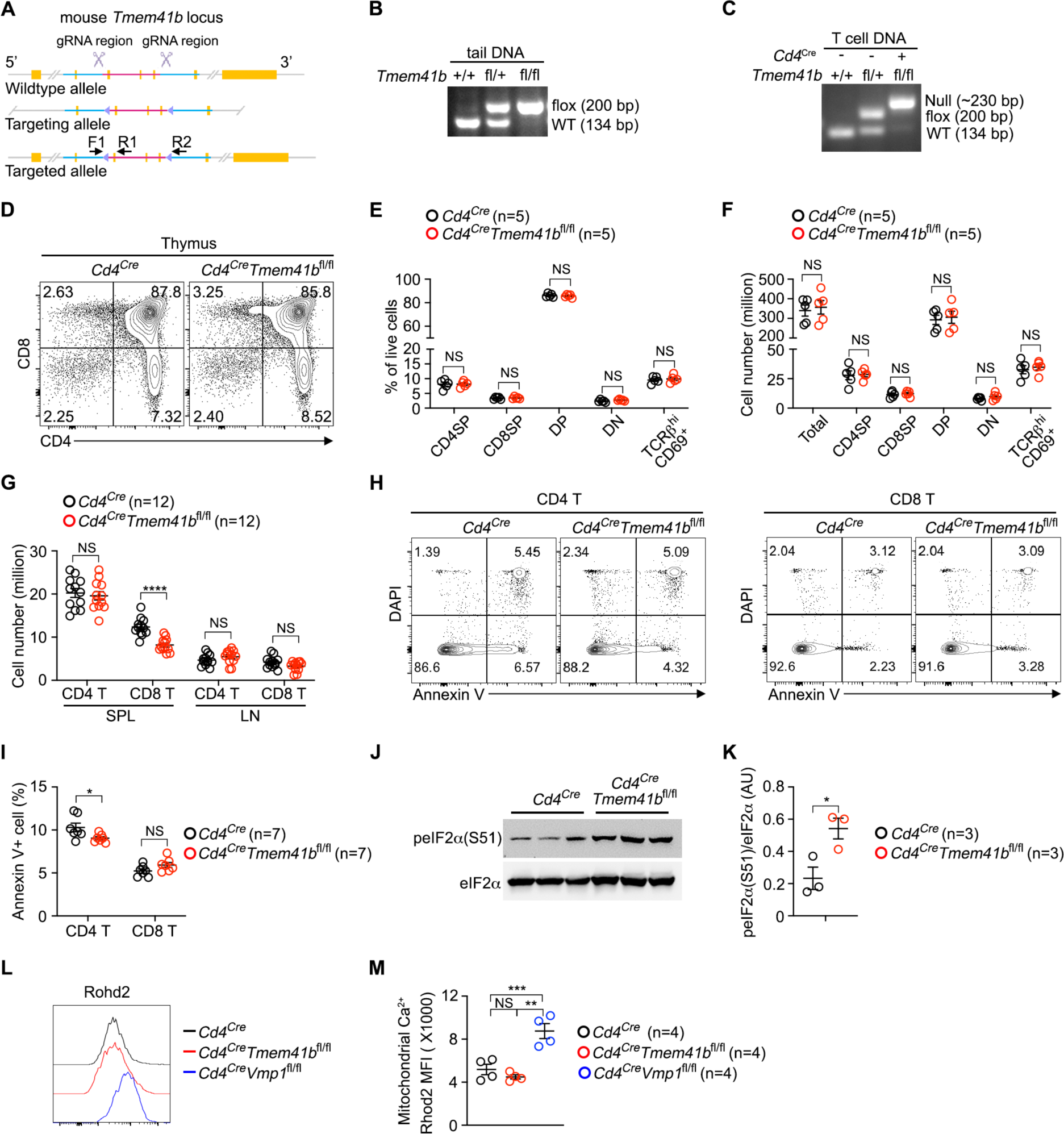
Analysis of T cell-specific TMEM41B knockout mice, related to Figure 1 and Figure 4. (A) Gene targeting strategy for generation of a *Tmem41b* flox allele. (B) Genotyping of DNA from tail of mice with indicated genotypes. (C) Genotyping of DNA from T cells of mice with indicated genotypes. (D - F) Flow cytometry analysis of thymocytes from control (*Cd4*^Cre^) and TMEM41B-deficient (*Cd4*^Cre^*Tmem41b*^fl/fl^) mice. Representative plots (D) and statistics (E and F) are shown. CD4^+^CD8^-^ (CD4SP), CD4^-^CD8^+^ (CD8SP), CD4^+^CD8^+^ (DP), CD4^-^CD8^-^ (DN). (n = 5 mice). (G) T cell number in spleen (SPL) and peripheral lymph nodes (pLN) of control and TMEM41B-deficient mice. (n = 12 mice). (H and I) Flow cytometry analysis of T cell apoptosis. Representative plots (H) and statistics (I) are shown. (n = 7 mice) (J and K) Immunoblot of eIF2α and its phosphorylation form (peIF2α-S51) in T cells freshly isolated from control and TMEM41B-deficient mice. Each lane represents an individual mouse. (n = 3 mice). (L and M) Flow cytometry analysis of mitochondrial Ca^2+^ in control and TMEM41B-deficient CD4 T cells. Representative plots (L) and statistics (M) are shown. (n = 4 mice). * p< 0.05, ** p< 0.01, *** p< 0.001, NS, not significant by two-tailed unpaired t-test (E, F, I, K) or one-way ANOVA (M).

**Figure S4.**
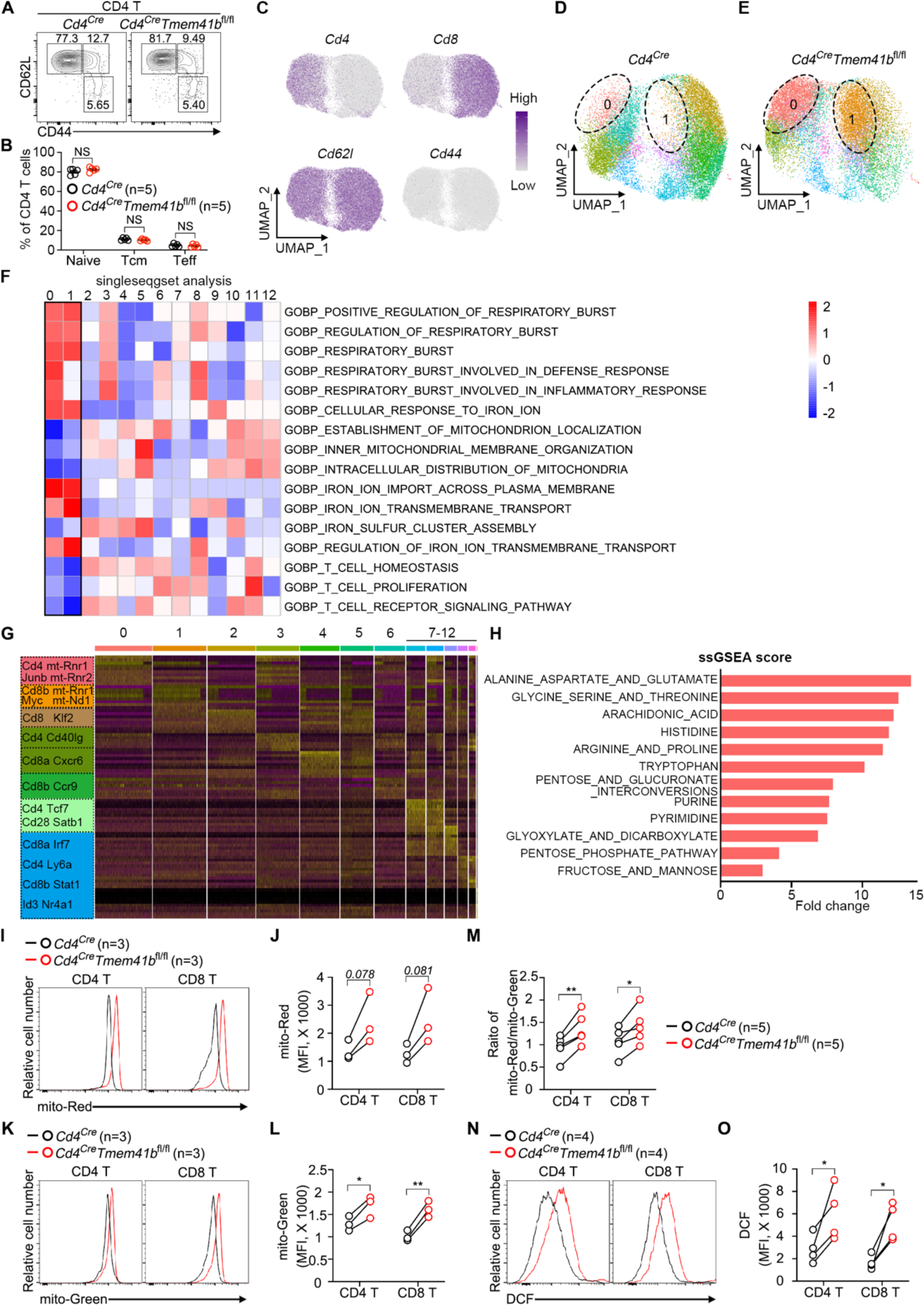
Metabolic activation of TMEM41B-deficient naive T cell, related to Figure 4. (A and B) Flow cytometry analysis of activation status (CD44 vs CD62L) of CD4 T cells from peripheral lymph nodes (pLN) of control (*Cd4*^Cre^) and TMEM41B-deficient (*Cd4*^Cre^*Tmem41b*^fl/fl^) mice (6 ∼ 8-week-old). Representative plots (A) and statistics (B) are shown. (n = 5 mice). (C) Uniform manifold approximation and projection (UMAP) plots of control and TMEM41B-deficient naive T cells. (D and E) UMAP plots of control (D) and TMEM41B-deficient (E) naive T cells, highlighting cells in cluster of 0 and 1. (F) Gene-set enrichment analysis (GSEA) of all clusters in naive T cells by singleseqgset analysis method. (G) Clustered heatmap of representative marker genes across clusters. (H) Metabolism-related GSEA of up-regulated genes in TMEM41B-deficient naive T cells versus control naive T cells by ssGSEA method. (I - M) Flow cytometry analysis of mitochondrial mass and mitochondrial membrane potential of naive T cells with indicated genotypes. Representative plots (I and K) and statistics (J, L and M) are shown. (n = 3 or 5 mice). (N and O) Flow cytometry analysis of reactive oxygen species (ROS) in control and TMEM41B-deficient naive T cells. Representative plots (N) and statistics (O) are shown. (n = 4 mice). * p< 0.05, ** p< 0.01, NS, not significant by two-tailed unpaired t-test (B) or two-tailed paired t-test (J, L, M and O).

**Figure S5.**
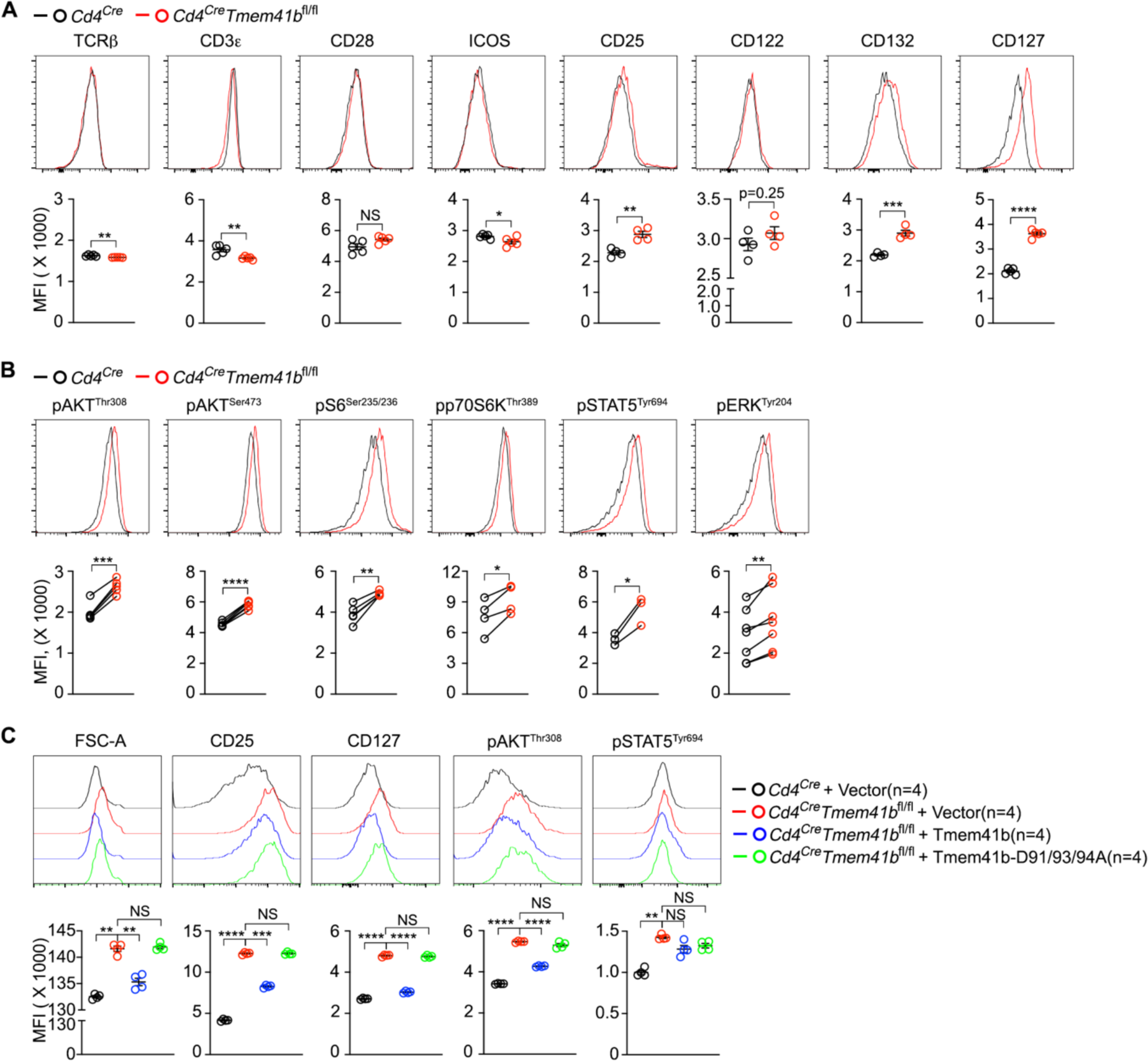
TMEM41B represses basal IL-2 and IL-7 receptor signaling in naive CD8 T cells via Ca^2+^ channel activity, related to Figure 5. (A) Flow cytometry analysis of the expression of indicated proteins on naive CD8 T cells from peripheral lymph nodes (pLN) of control (*Cd4*^Cre^) and TMEM41B-deficient (*Cd4*^Cre^*Tmem41b*^fl/fl^) mice. Representative plots and statistics are shown. (n = 4 or 5 mice). (B) Flow cytometry analysis of the phosphorylation of indicated proteins in naive CD8 T cells from pLN of control and TMEM41B-deficient mice. Representative plots and statistis are shown. (n = 3 ∼ 8 mice). (C) Activated CD8 T cells were transduced with indicated retroviral constructs. Cell size, CD25/CD127 expression and phosphorylation of AKT/STAT5 were measured by flow cytometry 72 hours post-transduction. Representative plots and statistics are shown. (n = 4 independent experiments). * p< 0.05, ** p< 0.01, *** p< 0.001, **** p< 0.0001, NS, not significant by two-tailed unpaired t-test (A), two-tailed paired t-test (B), or one-way ANOVA (C).

**Figure S6.**
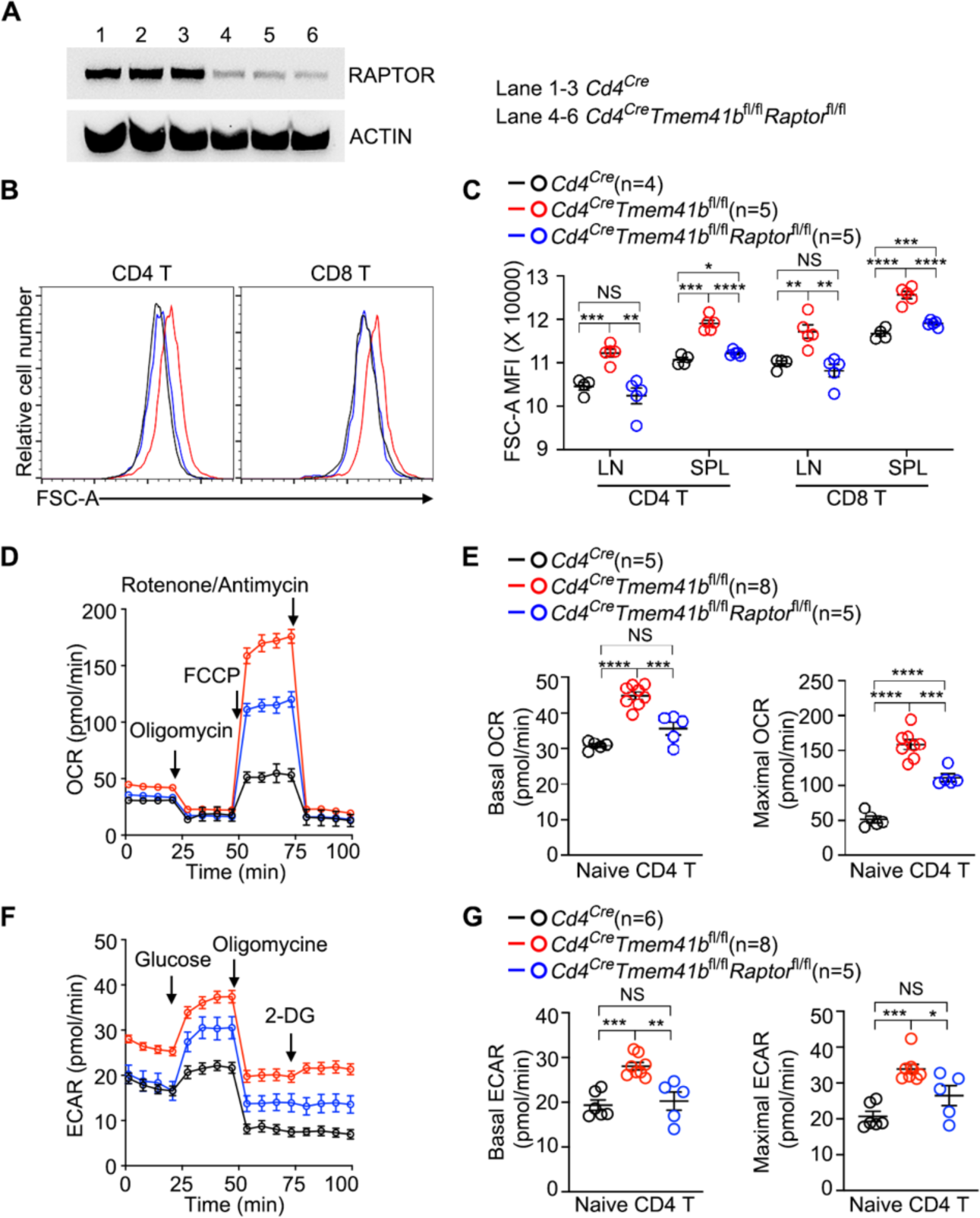
mTORC1 hyperactivation contributes to metabolic activation of TMEM41B-deficient naive T cells, related to Figure 5. (A) Immunoblot of RAPTOR in total T cells from control (*Cd4*^Cre^) and TMEM41B-deficient (*Cd4*^Cre^*Tmem41b*^fl/fl^) mice. Each lane represents an individual mouse. (B and C) Flow cytometry analysis of cell size of naive T cells from control and TMEM41B-deficient mice. Representative plots (B) and statistics (C) are shown. (n = 4 or 5 mice). (D and E) Oxygen consumption rate (OCR) of naive CD4 T cells from indicated genotypes were measured with a Mito stress test kit. The plots (D) and calculated basal and maximal OCR (E) are shown. (n = 5 or 8 independent experiments). (F and G) Extracellular acidification rate (ECAR) of naive CD4 T cells with indicated genotypes were measured with a Glycolysis stress test kit. The plots (F) and calculated basal and maximal ECAR (G) are shown. (n = 5, 6 or 8 independent experiments). * p< 0.05, ** p< 0.01, *** p< 0.001, **** p< 0.0001, NS, not significant by one-way ANOVA (C, E and G).

**Figure S7.**
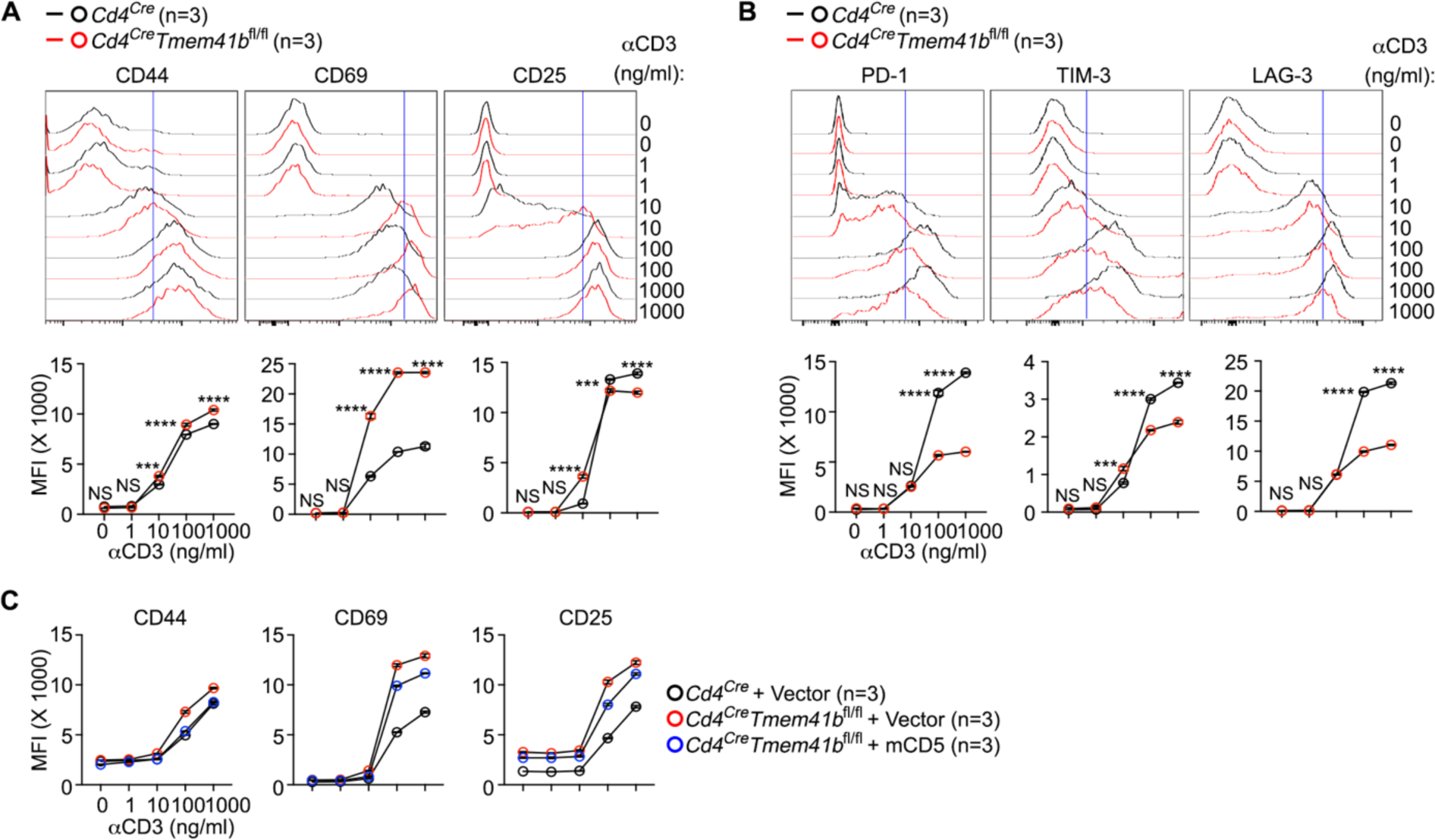
TMEM41B represses T cell responsiveness via CD5, related to Figure 6. (A) Flow cytometry analysis of the upregulation of CD44, CD49 and CD25 on CD8 T cells upon stimulation with different doses of αCD3 antibodies. Representative plots and statistics are shown. (n = 3 independent experiments). (B) Flow cytometry analysis of the upregulation of PD-1, TIM-3 and LAG-3 on CD8 T cells upon stimulation with different doses of αCD3 antibodies. Representative plots and statistics are shown. (n = 3 independent experiments). (C) Flow cytometry analysis of CD44, CD69 and CD25 expression on CD8 T cells transduced with indicated constructs upon αCD3 stimulation. (n = 3 independent experiments). *** p< 0.001, **** p< 0.0001, NS, not significant by two-way ANOVA (A and B).

